# *O*-GlcNAcylation, oxidation and CaMKII contribute to atrial fibrillation in type 1 and type 2 diabetes by distinct mechanisms

**DOI:** 10.1101/2020.02.18.954909

**Authors:** Olurotimi O. Mesubi, Adam G. Rokita, Neha Abrol, Yuejin Wu, Biyi Chen, Qinchuan Wang, Jonathan M. Granger, Anthony Tucker-Bartley, Elizabeth D. Luczak, Kevin R. Murphy, Priya Umapathi, Partha S. Banerjee, Tatiana N. Boronina, Robert N. Cole, Lars S. Maier, Xander H. Wehrens, Joel L. Pomerantz, Long-Sheng Song, Rexford S. Ahima, Gerald W. Hart, Natasha E. Zachara, Mark E. Anderson

## Abstract

Diabetes mellitus and atrial fibrillation (AF) are major unsolved public health problems, and diabetes is an independent risk factor for AF in patients. However, the mechanism(s) underlying this clinical association is unknown. Elevated protein *O*-GlcNAcylation (OGN) and reactive oxygen species (ROS) are increased in diabetic hearts, and calmodulin kinase II (CaMKII) is a proarrhythmic signal that may be activated by OGN (OGN-CaMKII) and ROS (ox-CaMKII). We induced type 1 (T1D) and type 2 diabetes (T2D) in a portfolio of genetic mouse models capable of dissecting the role of OGN and ROS at CaMKII and the type 2 ryanodine receptor (RyR2), an intracellular Ca^2+^ channel implicated as an important downstream mechanism of CaMKII-mediated arrhythmias. Here we show that T1D and T2D significantly increased AF, similar to observations in patients, and this increase required CaMKII. While T1D and T2D both require ox-CaMKII to increase AF, they respond differently to loss of OGN-CaMKII or OGN inhibition. Collectively, our data affirm CaMKII as a critical proarrhythmic signal in diabetic AF, and suggest ROS primarily promotes AF by ox-CaMKII, while OGN promotes AF by diverse mechanisms and targets, including CaMKII and RyR2. The proarrhythmic consequences of OGN- and ox-CaMKII differ between T1D and T2D. These results provide new and unanticipated insights into the mechanisms for increased AF in diabetes mellitus, and suggest successful future therapies will need to be different for AF in T1D and T2D.

## Introduction

Atrial fibrillation (AF) and diabetes mellitus (DM) are major, unsolved public health problems (1–3). The incidence and prevalence of both conditions are projected to increase significantly in the United States and worldwide in the coming decades (1, 4–7). AF is the most common clinical arrhythmia (8), and it is associated with significant morbidity and mortality (9–12), such as stroke and heart failure. It has become increasingly clear that DM, both type 1 DM (T1D) and type 2 DM (T2D), is an independent risk factor for AF (13–15), and the coexistence of AF and DM results in increased mortality, suffering, and cost (16–18). However, current therapies, such as antiarrhythmic drugs and catheter ablation (19, 20), are inadequate. Thus, improved understanding of the molecular mechanisms connecting DM and AF is an important goal for developing more effective therapies.

DM is characterized by hyperglycemia, elevated levels of reactive oxygen species (ROS) (21, 22) and increased *O*-GlcNAcylation (OGN) (23, 24). OGN is an evolutionarily conserved nutrient and stress sensing post-translational protein modification that occurs by covalent attachment of single O-linked *N*-acetylglucosamine (*O*-GlcNAc) residues to the amino acids serine and threonine, resulting in alteration of protein function akin to phosphorylation (25, 26). This dynamic process is controlled exclusively by two enzymes – *O*-GlcNAc transferase (OGT), which catalyzes the addition of *O*-GlcNAc to target residues, and *O*-GlcNAcase (OGA), which catalyzes the removal of *O*-GlcNAc (25, 26). Atrial myocardium from diabetic patients has increased ROS (27, 28) and protein OGN (29). Both ROS and OGN are associated with diabetic cardiomyopathy, and atrial fibrillation (21, 30–34), but it is uncertain if these changes contribute to proarrhythmic molecular pathways favoring atrial fibrillation (21, 35, 36).

The multifunctional calcium and calmodulin-dependent protein kinase II (CaMKII) has emerged as a proarrhythmic signal in AF in the absence of DM (32, 37, 38). Work by us and others, suggests that ROS (39) and OGN (31) can modify the CaMKII regulatory domain, causing CaMKII to become constitutively active, leading to inappropriate Ca^2+^ leak from myocardial type 2 ryanodine receptors (RyR2), triggered action potentials and arrhythmias (31, 32, 40–43). Intriguingly, the ROS (methionines 281/282, ox-CaMKII) and OGN (serine 280, OGN-CaMKII) modified residues of the CaMKII regulatory domain are adjacent (Fig 1A), suggesting the hypothesis that CaMKII may integrate these upstream signals – ROS and OGN, which are present in diabetic atrium to favor AF. The role of these signals, and whether they operate independently or together, is untested in vivo. A recent study implicates OGN-CaMKII in increased ROS production and CaMKII signaling independent of ox-CaMKII in isolated ventricular myocytes in response to acute hyperglycemia (44). However, the potential proarrhythmic contributions of ox-CaMKII and OGN-CaMKII have not been directly determined in vivo, in atria nor in diabetic AF.

**Figure 1.**
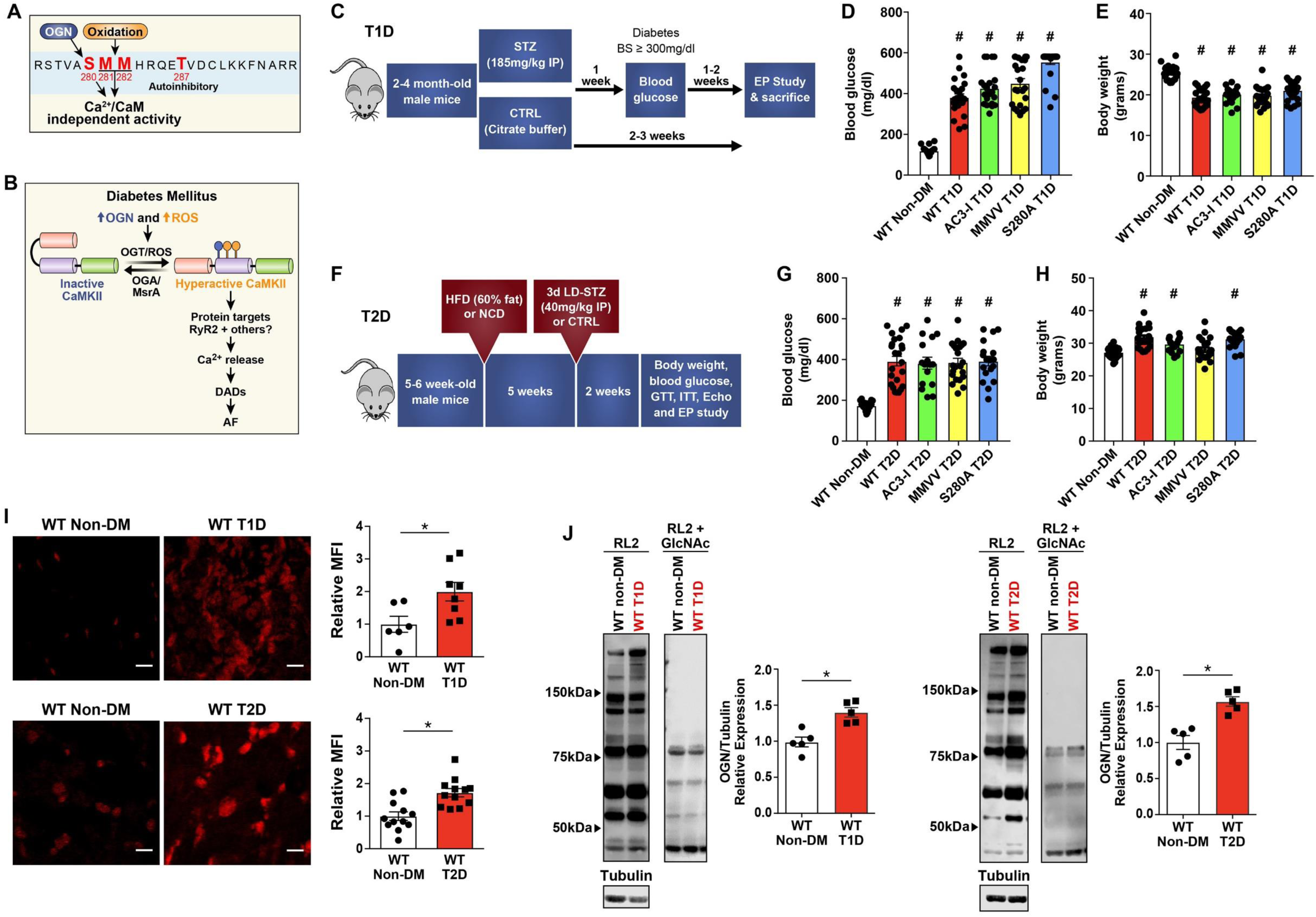
Reactive oxygen species (ROS) and O-GlcNAcylation (OGN), upstream activating signals for CaMKII, are elevated in type 1 (T1D) and type 2 (T2D) diabetic hearts. (A) Oxidation at methionines 281/282 and OGN at serine 280 are key post-translational modifications hypothesized to contribute to diabetic heart disease and arrhythmias. (B) Schematic representation of proposed hypothesis that excessive ROS and OGN in DM promotes AF through CaMKII dependent signaling. (C) Schematic diagram of diabetes induction and experimental protocol for T1D. (D) Elevated blood glucose in STZ treated (T1D) mice compared to non-diabetic citrate buffer treated (non-DM) control mice at the time of electrophysiology study, two weeks after STZ injection. (E) Summary data for body weight two weeks after STZ injection. (F) Schematic diagram of diabetes induction and experimental protocol for T2D. (G) Elevated blood glucose in T2D mice compared to non-diabetic controls for T2D at the time of electrophysiology study, two weeks after STZ injection. (H) Summary data for body weight two weeks after LD-STZ injection. (I) Representative confocal images (original magnification, x40) and summary data for DHE fluorescence in mouse atrial tissue show increased ROS in T1D (top panel) and T2D (bottom panel). Scale bars 10µm (n = 6 WT non-DM, n = 8 WT T1D, n = 4 WT non-DM, n = 4 WT T2D) (J) Representative western blots for total OGN modified protein levels (OGN monoclonal antibody – RL2) and competition assay (RL2 + 500mM GlcNAc) from heart lysates, and summary data for total OGN modified protein levels normalized to tubulin from heart lysates from T1D (left panel) and T2D (right panel) (n = 3 – 7/group). OGN protein quantification excluded the non-competed bands. DADs, delayed after depolarizations; DHE, dihydroethidium; HFD, high fat diet; LD-STZ, low dose STZ; MFI, mean fluorescent intensity; NCD, normal chow diet; OGN, O-GlcNAcylation; ROS, reactive oxygen species; RyR2 – Ryanodine receptor type 2; STZ, streptozocin; WT, wild type. Data are represented as mean ± s.e.m, unless otherwise noted. Statistical comparisons were performed using two-tailed Student’s t test or one way AVOVA with Tukey’s multiple comparisons test for continuous variables and Fischer’s exact test for dichotomous variables. (#p < 0.0001 versus WT non-DM, *p < 0.05 versus WT non-DM).

We reasoned that hyperglycemia secondary to DM is proarrhythmic and promotes AF through increased ROS and OGN upstream of CaMKII with consequent activation of RyR2, and potentially other targets, leading to increased proarrhythmic triggered activity (Fig 1B). We performed new experiments to investigate the molecular mechanism(s) by which CaMKII integrates upstream signals from ROS and OGN to promote AF in DM, to understand if CaMKII activation by ROS and/or OGN are critical proarrhythmic mechanisms for diabetic AF, and if the proarrhythmic consequences of CaMKII are similar or different in T1D and T2D. We induced T1D or T2D in a panel of new and established mouse models where ROS and OGN activation of CaMKII was selectively ablated, where ROS and OGN upstream pathways were controlled, and where convergent actions of CaMKII on RyR2 were prevented or mimicked. Our results establish CaMKII as a critical sensor of ROS and hyperglycemia, and CaMKII activation of RyR2 as an important proarrhythmic pathway for AF in DM. We found AF in T1D and T2D required ox-CaMKII. However, AF induction in T1D and T2D exhibited different responses to loss, by knock in replacement, of a CaMKII OGN site (S280A), or transgenic myocardial over-expression of OGA. We discovered that transgenic myocardial OGA over-expression protected against diabetic AF by CaMKII-dependent and -independent pathways.

## Results

### T1D and T2D mice have increased ROS and OGN

To test the hypothesis that CaMKII contributes to hyperglycemia enhanced AF susceptibility, our first approach was to use suitable mouse models of T1D and T2D and determine myocardial levels of ROS and OGN. For the T1D mouse model, we injected mice with a single dose (185 mg/kg, i.p.) of streptozocin (STZ) (Fig 1C) (45, 46). STZ treated (T1D) mice had significantly higher blood glucose (Fig 1D and Supplementary Fig 1A) and reduced body weight (Fig 1E and Supplementary Fig 1B) compared with placebo (citrate buffer) treated mice. There was no difference in heart weight indexed for body weight (Supplementary Fig 1C). T1D mice had slower heart rates (Supplementary Fig 1D), consistent with reports of defective heart rate in diabetic mice (46) and patients (47, 48). Echocardiographic measurements showed impaired left ventricular (LV) systolic function in the T1D mice, compared with non-diabetic littermate controls (Supplementary Fig 1E) without LV hypertrophy or dilation (Supplementary Fig 1F - H), findings similar to patients with diabetic cardiomyopathy (49). The T1D mice showed no evidence of ketoacidosis or renal failure (Supplementary Table 1). Thus, the T1D mice showed marked elevation in glucose, weight loss, and modest, but consistent, impairment in myocardial function, similar to previous reports of diabetic cardiomyopathy (50).

We considered the T1D model an ideal experimental platform to test the direct link between hyperglycemia and AF. However, the predominant form of DM in the human adult population affected by AF is T2D (51), a more complex disease characterized by hyperglycemia, impaired insulin secretion and insulin resistance (52). Thus, we next generated a T2D mouse model using five- to six-week-old male C57BL/6J mice fed a high-fat diet; five weeks after initiation of high fat diet, the mice received daily injections of low dose STZ (40mg/kg/day, i.p.) for three consecutive days (Fig 1F). The T2D mice had elevated blood glucose (Fig 1G), increased body weight, albeit with a slight decrease after STZ injection (Fig 1H and Supplementary Fig 2A), no change in fasting insulin (Supplementary Fig 2B), but increased insulin resistance, determined by the homeostatic model assessment of insulin resistance (HOMA-IR) (53) (Supplementary Fig 2C), compared to untreated wild type (WT) controls. Consistent with the defining features of T2D, our mice showed reduced glucose and insulin tolerance (Supplementary Fig 2D and 2E) two weeks after low dose STZ injection.

We next measured atrial ROS and OGN from whole heart lysates using from WT T1D, T2D and non-diabetic control hearts. Consistent with prior animal (21, 22, 30, 34) and human data (27, 28, 34), T1D and T2D heart tissues showed increased ROS (Fig 1I) and OGN (Fig 1J) compared to WT non-diabetic controls. These foundational data suggested these models would be useful for testing the hypothesis that ox-CaMKII and OGN-CaMKII contributed to AF in T1D and T2D.

### Increased AF susceptibility in T1D and T2D is CaMKII-dependent

Excessive CaMKII activity is implicated in non-diabetic AF in patients (32, 37, 38), large animal models (37, 54, 55), and in mice (32, 37, 56). More recently, OGN-CaMKII has been linked to ventricular arrhythmias in diabetic cardiomyopathy (31, 44). To test if these diabetic mouse models are susceptible to increased AF, we performed rapid right atrial burst pacing in anesthetized mice using an established protocol (Fig 2A) (32, 37, 57), with modification of the definition of AF in this study (see Methods section). T1D mice had significantly higher AF compared to non-diabetic controls (Fig 2B). To further probe the effect of hyperglycemia on AF susceptibility, T1D mice were treated with insulin (delivered via osmotic pumps - LinBit) for one week after STZ injection. Insulin treated T1D mice with rescue of hyperglycemia (< 300 mg/dl, Fig 2C) had significantly reduced AF susceptibility compared to insulin treated T1D mice with persistent hyperglycemia (> 300 mg/dl, Fig 2D). These findings, taken together with other reports (14, 15, 58), confirmed that T1D is an AF risk factor in mice, similar to the situation in patients.

**Figure 2.**
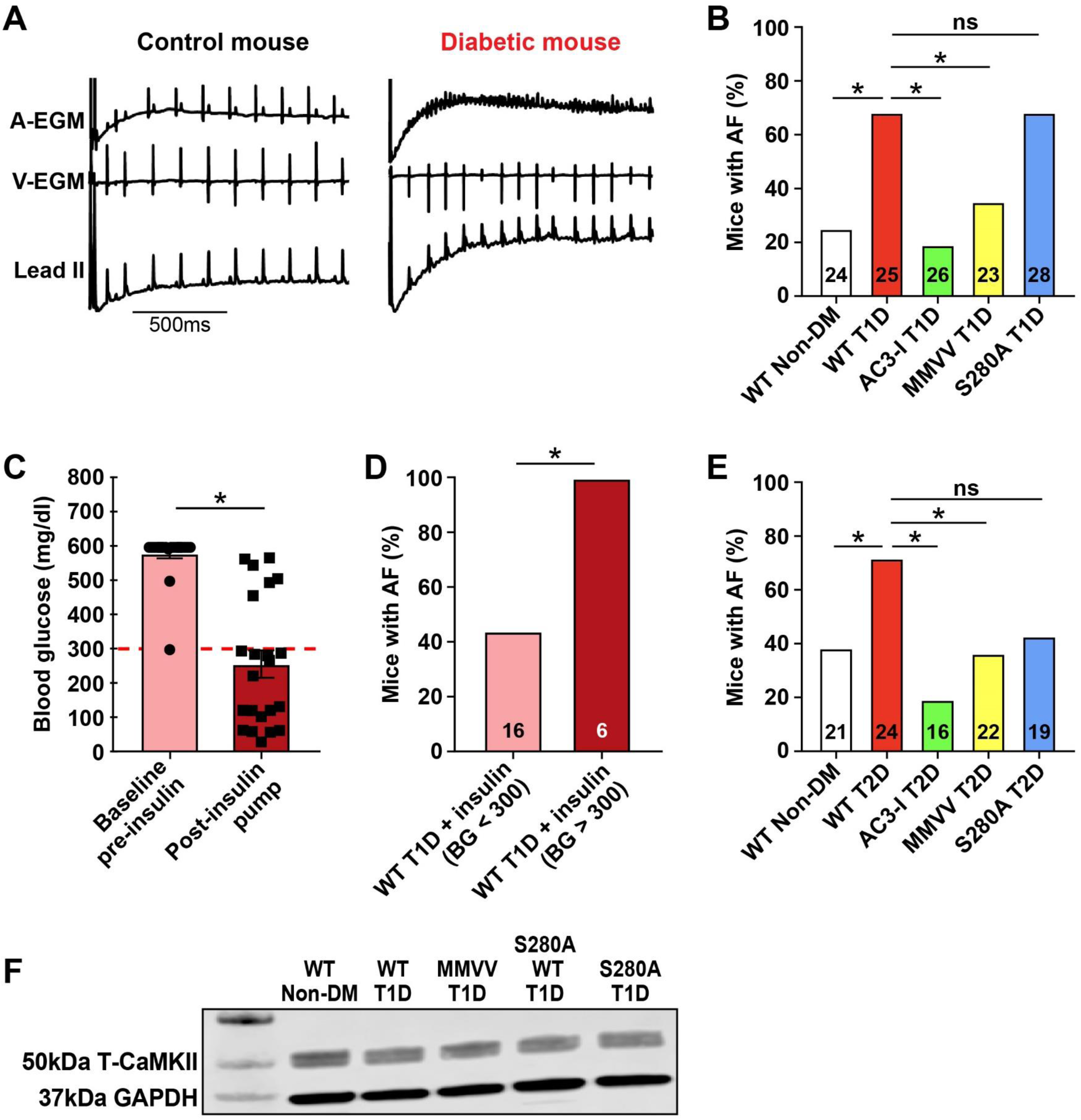
CaMKII promotes enhanced atrial fibrillation (AF) susceptibility in T1D and T2D diabetes mellitus. (A) Representative tracings of intra-cardiac (atrial, A-EGM; ventricular, V-EGM) and lead II surface electrocardiograms recorded immediately following rapid atrial burst pacing demonstrating normal sinus rhythm in a control non-DM wild type (WT) mouse and irregularly irregular atrial and ventricular electrical impulses marking AF in a diabetic WT T1D mouse. (B) Marked AF susceptibility in WT T1D mice compared to WT non-DM mice. This is reversed in AC3-I and MMVV T1D mice, but not in S280A T1D mice. (C) Pre- and post-insulin pump (LinBit) implantation blood glucose levels one week after STZ treatment (pre-insulin) and one week after insulin pump implantation (post-insulin). (D) Insulin treatment prevents enhanced AF in WT T1D mice with blood glucose (BG) level less than 300mg/dl on insulin treatment. (E) Increased AF susceptibility is present in WT T2D mice compared to non-diabetic controls; AC3-I and MMVV T2D mice are protected from enhanced AF and there is a trend towards protection in S280A T2D mice (p = 0.07). (F) Representative western blot for CaMKII and GAPDH in atrial lysates from WT non-DM and T1D WT, MMVV, S280A WT and S280A mice. EGM, electrogram; ns, non-significant. Data are represented as mean ± s.e.m, unless otherwise noted. The numerals in the bars represent the sample size in each group. Statistical comparisons were performed using two-tailed Student’s t test or one way AVOVA with Tukey’s multiple comparisons test for continuous variables and Fischer’s exact test for dichotomous variables. (*p <0.05).

To test if CaMKII contributes to hyperglycemia enhanced AF susceptibility, we used mice with myocardial targeted transgenic expression of AC3-I, a CaMKII inhibitory peptide (59). We found that AC3-I mice with T1D were protected from AF compared to WT non-diabetic controls (Fig 2B). Despite the reduction in AF, T1D AC3-I mice had similar increases in blood glucose as the WT T1D mice (Fig 1D). These findings supported the hypothesis that CaMKII is a proarrhythmic signal coupling DM to increased risk of AF. We next tested AF susceptibility in the T2D mouse model (Fig 1F). The T2D mice had significantly higher AF compared to non-diabetic controls (Fig 2E), similar to the findings in the T1D mice and in humans with T1D and T2D (14, 15). To determine if CaMKII contributes to AF susceptibility in T2D, we performed right atrial burst pacing in AC3-I T2D mice. Similar to our findings in T1D, AC3-I T2D mice were protected from AF (Fig 2E) compared with non-diabetic controls, and had similar levels of hyperglycemia as WT T2D mice (Fig 1G). Taken together, these findings support the hypothesis that CaMKII is a proarrhythmic signal that is essential for increased risk of AF in validated mouse models of T1D and T2D.

### Differential effect of MM281/282 and S280 on AF susceptibility in T1D and T2D

After initial activation, ROS and OGN can modify the CaMKII regulatory domain to lock CaMKII into a constitutively active, Ca^2+^ and calmodulin independent conformation that is established to promote arrhythmias and cardiomyopathy (60, 61). ROS activates CaMKII by oxidizing a pair of methionines (M281/282) (39), while OGN activates CaMKII by modifying serine 280 (S280) (31) (Fig 1A). The numbering is for CaMKIIδ, the most abundant CaMKII isoform in ventricular myocardium (62–64). We used qRT-PCR to measure expression of each CaMKII isoform in human and mouse atrium. We found that, similar to the situation in ventricle, that CaMKIIδ has the highest mRNA expression in human (Supplementary Fig 3A, Supplementary Table 2) and mouse atrium (Supplementary Fig 3B). In addition, CaMKII isoform mRNA expression profile was unchanged in T1D and T2D atria compared with WT non-diabetic controls (Supplementary Figs 3C and 3D).

To determine the role of ox-CaMKII and OGN-CaMKII in hyperglycemia primed-AF, we utilized CaMKIIδ genetically modified (knock-in) mice. Our lab previously generated an ox-CaMKII resistant (MMVV) knock-in mouse (46). These MMVV mice are notable for exhibiting resistance to sudden death due to severe bradycardia after myocardial infarction during T1D (46), and are protected from angiotensin II infusion primed AF (32). These findings provided support for the role of ox-CaMKII as a pathological signal in atrial myocardium. OGN activation of CaMKII occurs by modification of a S280 (31), so we developed an OGN-CaMKII resistant (S280A) mouse using CRISPR/Cas9 (65–67) (Supplementary Fig 4A). S280A mice were born in Mendelian ratios, had normal morphology and similar baseline left ventricular (LV) fractional shortening (Supplementary Fig 4B) and LV mass (Supplementary Fig 4C), compared to WT littermate controls. MMVV, S280A, and WT mice have similar total myocardial CaMKII expression (Fig 2F), and resting heart rates (Supplementary Fig 4D). We interpreted these findings, taken together with previous studies on MMVV mice (32, 46), to suggest that the S280 and M281/282 sites on CaMKIIδ are dispensable for normal development and basal function.

We induced T1D in MMVV and S280A mice, and performed AF induction studies. T1D MMVV mice were resistant to AF, but T1D S280A mice showed similar AF susceptibility as T1D WT mice (Fig 2B). The WT and S280A T1D mice showed a modest, but consistent decrement in LV fractional shortening compared to non-diabetic counterparts (Supplementary Fig 1E and 4F). In contrast, MMVV mice were resistant to T1D induced LV dysfunction (Supplementary Fig 4F). The T1D MMVVV and S280A mice demonstrated similar increases in blood glucose, and comparable body weight (Fig 1D and 1E; Supplementary Fig 4G and 4H). These results suggested that ox-CaMKII at MM281/282, but not OGN-CaMKII at S280, contributed to T1D cardiomyopathy and AF. We next performed AF induction studies in T2D MMVV and S280A mice (Fig 2E). T2D MMVV mice were resistant to AF, while T2D S280A mice had a trend towards rescue (p = 0.07). We interpret these data to suggest that ox-CaMKII contributes to hyperglycemia primed-AF in both T1D and T2D. In contrast, the S280 site on CaMKIIδ contributes to AF in T2D but not in T1D.

### Attenuation of ROS signaling prevents AF due to hyperglycemia

In contrast to the differential response of T1D and T2D to manipulation of OGN and S280A for AF prevention, data on the role of M281/282 for promoting AF were highly consistent in both diabetic models. We next focused on the T1D model, because of its relative simplicity, to test for cellular sources of ROS that were important for ox-CaMKII and AF. Major ROS sources in heart include mitochondria and NADPH oxidases. We used a combination of genetic and pharmacologic approaches to inhibit ROS. Based on our earlier discovery that mitochondrial ROS increases ox-CaMKII in vitro during hyperglycemia (46), we tested if mitochondrial antioxidant therapy could protect against AF in T1D. We found that mice treated with daily injections of MitoTEMPO (1mg/kg, i.p.), a mitochondrial targeted antioxidant (68), for seven days, starting one week after STZ injection, were protected from AF in T1D (Fig 3A). In contrast, T1D mice treated with triphenylphosphonium (TPP), the mitochondrial targeting moiety of MitoTEMPO that lacks the anti-oxidant TEMPO, were not protected from AF (Fig 3A). MitoTEMPO and TPP treated mice had similar levels of hyperglycemia (Fig 3B). We confirmed increased mitochondrial ROS in atrial myocytes isolated from T1D using MitoSOX (Fig 3C), a fluorescent probe used for detection of mitochondrial ROS. NADPH oxidase is another important source of extra-mitochondrial ROS for increasing ox-CaMKII and promoting AF (31, 32, 69). Mice with T1D lacking p47 (*p47^-/-^* mice), an important component of myocardial NADPH oxidases (32, 39, 70), were also protected from AF (Fig 3A) and had similar levels of hyperglycemia as WT T1D mice (Fig 3B). Taken together, these data suggested that reduction of ROS signaling either by NADPH oxidase or mitochondrial ROS in the setting of T1D was sufficient to reduce AF risk.

**Figure 3.**
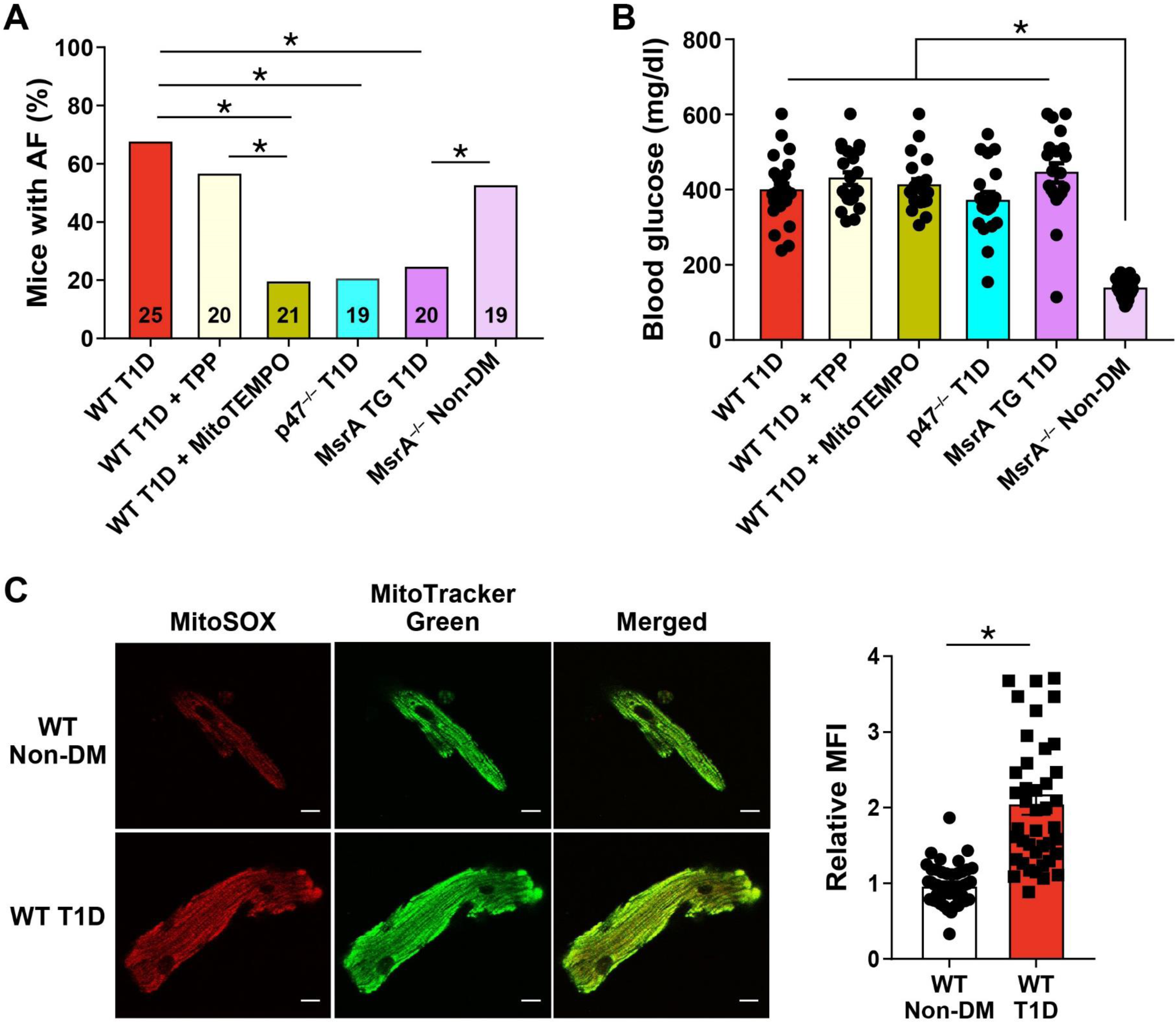
Targeted ROS inhibition and MsrA overexpression protects against AF in diabetes. (A) Inhibition of mitochondrial ROS by MitoTEMPO treatment, or inhibition of cytoplasmic ROS by loss of the p47 subunit of NADPH oxidase (p47^-/-^ mice) protect against AF in T1D mice. Mice with myocardial targeted transgenic overexpression of methionine sulfoxide reductase A (MsrA TG) were protected from T1D primed AF, while non-diabetic mice lacking MsrA (MrsA^-/-^ non-DM) showed increased AF susceptibility in the absence of diabetes. (B) Summary data of blood glucose measurements at the time of electrophysiology study. (C) Increased mitochondrial ROS in isolated atrial myocytes from WT T1D (lower panel) compared to WT non-DM (upper panel) mice detected by MitoSOX fluorescence. Representative confocal fluorescent images (original magnificent, x40) show MitoSOX (red, left), MitoTracker Green (green, middle) and merged images (right). Scale bars 10µm. (n = 41 – 47 cells in each group from 2 mice per group). Data are means ± s.e.m., and comparisons were evaluated by Student’s two-tailed t test (*p < 0.05).

Ox-CaMKII can be reduced by methionine sulfoxide reductase A (MsrA), and mice lacking MsrA (*MsrA^-/-^*) show increased ox-CaMKII (39). In contrast, mice with transgenic myocardial overexpression of MsrA are protected from ox-CaMKII induced cardiomyopathy (71). We found that non-diabetic *MsrA^-/-^* mice had easily induced AF, similar to WT mice with T1D (Fig 3A), and that MsrA transgenic mice with T1D were protected against AF, similar to non-diabetic WT mice (Fig 3A). The antiarrhythmic responses to mitoTEMPO infusion or MsrA over-expression were not related to effects on hyperglycemia because blood glucose levels were similarly increased in all mice (Fig 3B). We interpreted these findings as an indication that methionine oxidation and ox-CaMKII are elements of a conserved proarrhythmic pathway downstream to ROS in T1D. Further, decreasing methionine reductive capacity is sufficient to increase AF in the absence of DM.

### Hyperglycemia activates CaMKII in an MM281/282 ox-CaMKII but not S280 OGN-CaMKII dependent manner

The findings in the S280A mice in T1D were unexpected because of published evidence that OGN of CaMKII at S280 is required for hyperglycemia induced ROS, SR Ca^2+^ leak and triggered ventricular arrhythmias (31, 44). In order to determine if increased OGN detected in diabetic mouse myocardium could directly increase CaMKII activity, we used a novel fluorescent kinase reporter, recently validated in RPE-1 cells and mouse skeletal muscle (72, 73), based on work by Regot and colleagues (72). The fluorescent kinase translocation reporter (KTR) dynamically moves from the nucleus to the cytosol in response to kinase activation (Fig 4A) and the ratio of cytosol/nuclear distribution of the CaMKII-KTR fluorescent signal is a reliable measure of intracellular CaMKII activity (73). Isolated neonatal mouse cardiomyocytes transfected with CaMKII-KTR virus, 24 hours of after isolation, were cultured in low glucose (5.5 mM glucose) or high glucose (30 mM glucose) for 18 hours. Incubation with high glucose resulted in increased CaMKII activation compared with low glucose conditions (Fig 4B and 4C). There was no increase in CaMKII activation in neonatal cardiomyocytes incubated in hyperosmotic, euglycemic conditions (5.5 mM glucose + 24.5 mM mannitol) compared to control glucose conditions. To determine if CaMKII is critical for the KTR changes under hyperglycemic conditions, we co-incubated neonatal cardiomyocytes with high (30 mM) glucose and a high affinity ATP-competitive CaMKII inhibitor – AS105 (74). AS105 prevented increased CaMKII activity in response to hyperglycemia. These findings were consistent with the notion that hyperglycemia activates CaMKII. As a further proof of concept, we incubated WT neonatal cardiomyocytes with caffeine (10 mM) – a CaMKII activator, for 20 minutes in the absence of hyperglycemia and observed a similar increase in CaMKII activity compared to high glucose (Fig 4B and 4C). Next, to assess the contribution of S280 and MM281/282 (Fig 1A) to CaMKII activation by hyperglycemia, we incubated neonatal cardiomyocytes isolated from CaMKIIδ MMVV and S280A pups in high glucose. MMVV knock-in prevented increased CaMKII activation in response to hyperglycemia while the S280A cardiomyocytes exhibited increased CaMKII activity in response to hyperglycemia (Fig 4B and 4C). Given the apparent importance of ROS and MM281/282 in activating CaMKII under hyperglycemic conditions, we co-incubated WT neonatal cardiomyocytes with high glucose and *N*-acetyl cysteine (NAC). Treatment with 2 mM NAC prevented CaMKII activation in response to hyperglycemia. Taken together, these findings demonstrate that CaMKII activation by hyperglycemia requires MM281/282 (ox-CaMKII) and ROS in neonatal cardiomyocytes.

**Figure 4.**
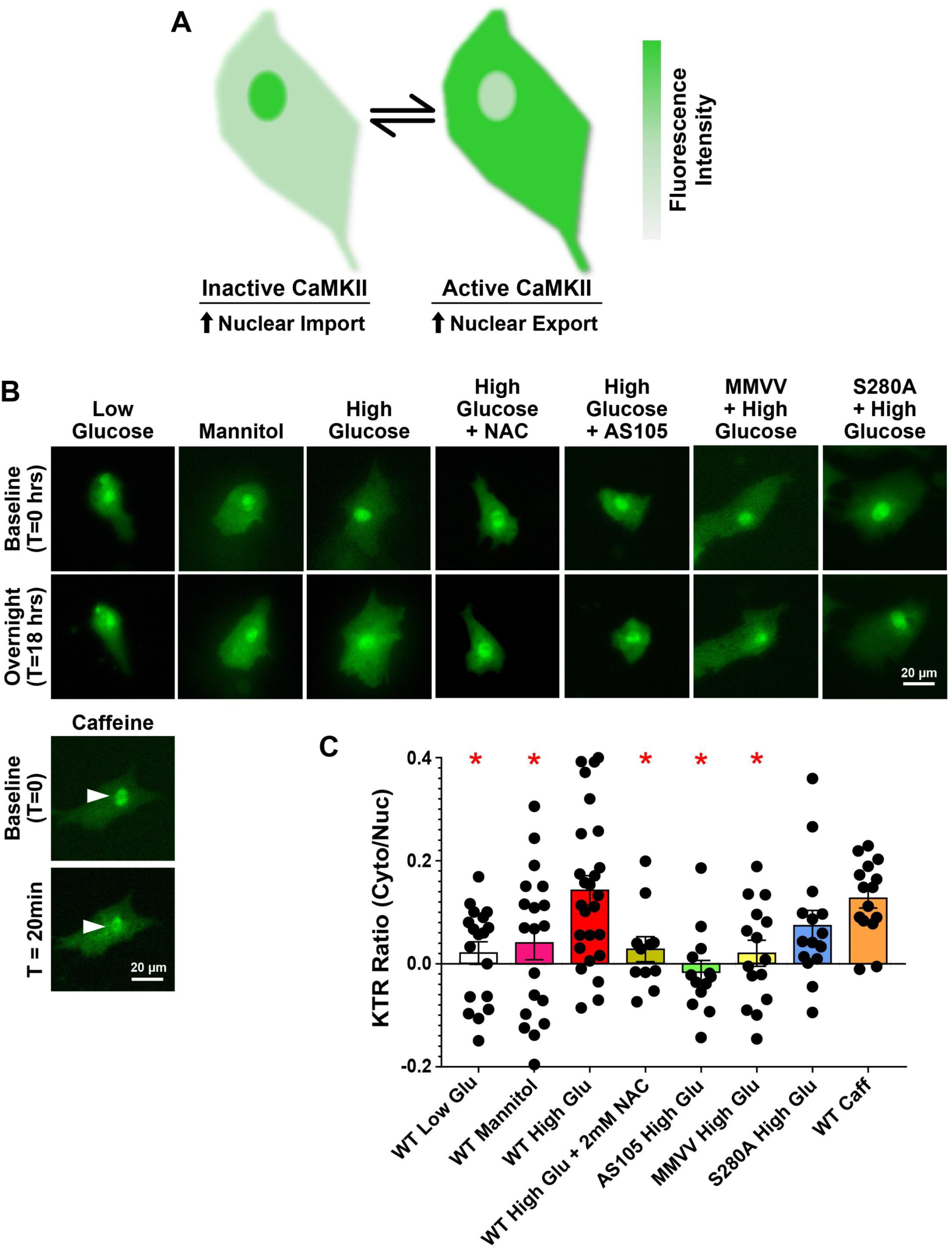
MM281/282 but not S280 is critical for CaMKII activation in response to hyperglycemia. (A) Schematic of CaMKII Kinase Translocation Reporter (CaMKII-KTR) Assay. CaMKII-KTR traffics between the nucleus and cytoplasm and phosphorylation by CaMKII results in net translocation of the KTR to the cytosol. Cytosolic to nuclear fluorescent signal ratio is a measure of CaMKII activity. (B) Representative fluorescent micrographs of KTR transfected neonatal mouse cardiomyocytes at baseline and time, t = 18 hrs post treatment. Cells from WT pups were incubated as indicated with 5.5 mM glucose (low glucose), 5.5 mM glucose + 24.5 mM mannitol (mannitol), 30 mM glucose (high glucose), high glucose + 2 mM N-acetyl cysteine (high glucose + NAC), or high glucose + 1 μM AS105 (a CaMKII inhibitor). Cells from MMVV and S280A pups were incubated with high glucose. Cells from WT pups at baseline and time, t = 20 minutes after treatment with 10 mM caffeine. Arrowheads indicate nuclei. (C) Summary data of the change in KTR cytosolic/nuclear ratio before and after treatments. Data are represented as mean ± s.e.m, and statistical comparisons were performed using one way ANOVA with Dunnett’s multiple comparisons test. (*p <0.05).

To further resolve OGN modification at S280 on CaMKII under euglycemic and hyperglycemic conditions, we performed targeted mass spectrometry analysis on CaMKII gel bands and immunoprecipitates from heart lysates from WT non-diabetic, T1D and T2D mice. Specifically, we focused on the region of interest in the regulatory domain of the CaMKII peptide with the amino acid sequence – STVASMMHR (Fig 1A and 1B), with several candidate posttranslational modification sites. We reliably detected two isoforms of CaMKII – γ and δ, with good coverage (46 to 58%) in the heart samples (Supplementary Table 3). We did not detect OGN modification at S280, despite several attempts using both data dependent and targeted mass spectrometry methods. In addition, we did not detect OGN modification at other sites on CaMKII. We were, however, able to detect HexNAc sugar modification on oxonium fragment ions on a different protein in these CaMKII enriched samples (data not shown). Interestingly, we detected phosphorylation at S280 on CaMKIIγ isoform in T1D hearts and on both CaMKIIδ (the predominant isoform in the heart) and CaMKIIγ isoforms in T2D hearts (Supplementary Fig 5A and 5B; and Supplementary Table 3 and 4). In confirmation of prior studies on CaMKII posttranslational modification, we identified oxidation at the paired methionines at MM281/282 and phosphorylation at T287, the first identified CaMKII posttranslational modification (75) (Supplementary Fig 5A – 5E; and Supplementary Table 3 - 5). We detected additional potential phosphorylation sites at S276 or T277 on the CaMKIIγ isoform in T2D hearts (Supplementary Fig 5C; and Supplementary Table 3 and 5). Although these findings do not rule out OGN modification at S280, our inability to detect this post-translational modification is consistent with our findings that S280A mice with T1D were not protected from AF. The identification of phosphorylation at the S280 on the CaMKIIδ isoform in T2D hearts is an interesting finding that may explain the partial rescue in S280A mice with T2D. Further, these findings suggest that effects of hyperglycemia via OGN modification are independent of CaMKII.

### OGN inhibition has a differential effect on AF susceptibility in T1D and T2D

Our studies up to this point did not address the possibility that increased OGN in T1D and T2D (Fig 1I) is proarrhythmic, potentially by actions at targets other than CaMKIIδ. As a first step we asked if preventing the synthesis of the *O*-GlcNAc transferase substrate UDP-GlcNAc, with the glutamine-fructose amidotransferase antagonist Diazo-5-oxonorleucine (DON), could reduce or prevent AF in diabetic mice. We administered DON (5mg/kg, i.p. injection) 30 minutes prior to AF induction. DON treatment significantly reduced AF susceptibility in WT T1D mice and S280A T1D mice (Fig 5A). MMVV T1D mice treated with DON had no significant further decrease in AF susceptibility (Fig 5A). These findings suggested that OGN contributed to diabetic AF in the T1D mouse model, but that this was independent of the S280 site on CaMKIIδ. We next asked if OGN inhibition had antiarrhythmic actions in T2D by treating T2D mice with DON prior to AF induction, as in the earlier T1D studies. Surprisingly, DON treatment was not protective against AF in T2D (Fig 5B). We interpreted these data up to this point to highlight important, but unanticipated differences in the AF mechanisms in T1D and T2D, and to suggest that separate antiarrhythmic therapies may be required for T1D and T2D, at least in mice.

**Figure 5.**
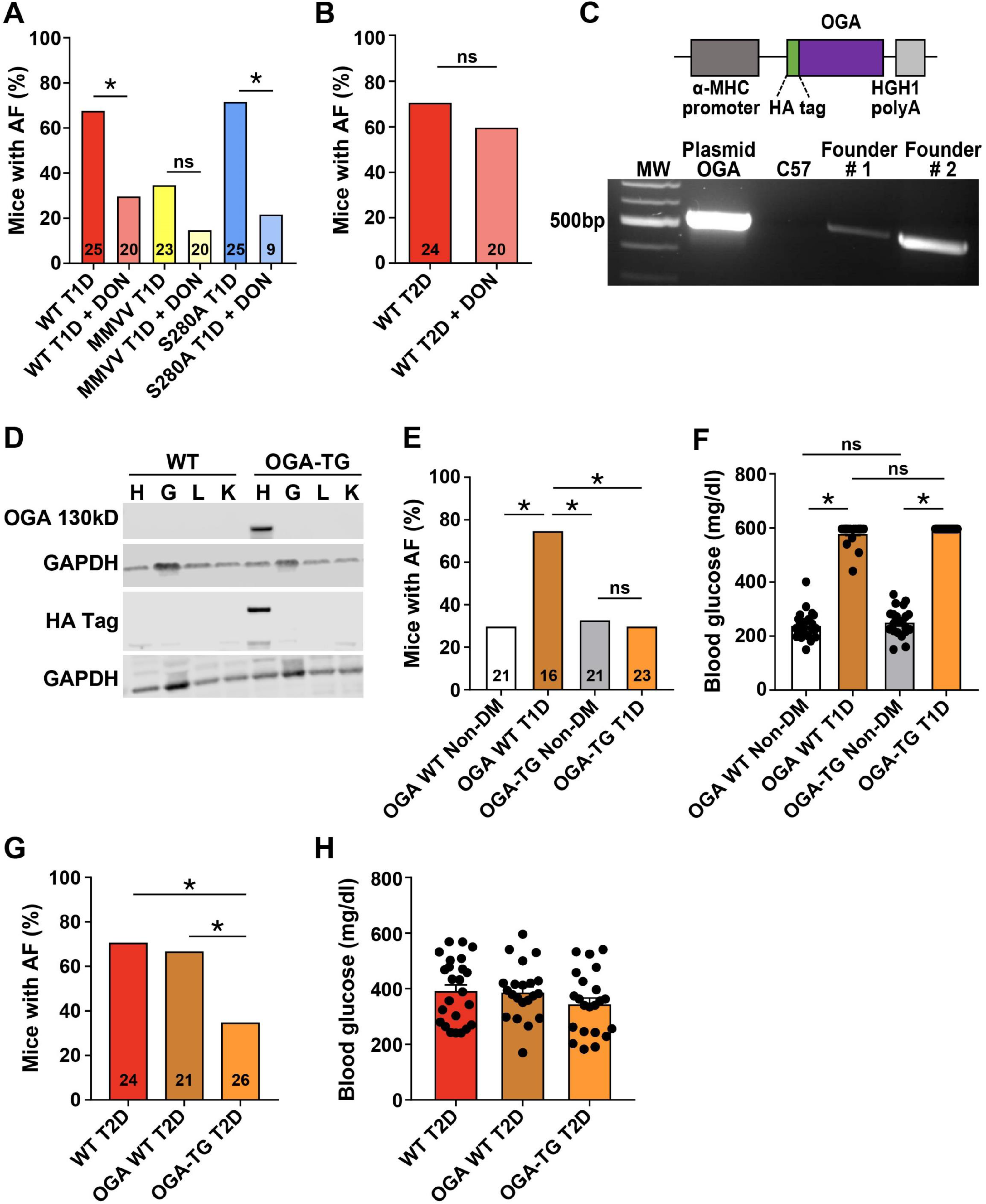
Differential responses of AF to DON and OGA in T1D and T2D. (A) DON pretreatment (5mg/kg i.p.) protected from AF in T1D WT and S280A mice, with no additional protection in T1D MMVV mice. (B) DON pretreatment did not protect from AF in T2D WT mice. (C) Schematic of the OGA (O-GlcNAcase) transgene construct with the α-myosin heavy chain (α-MHC) promoter, HA epitope marker and human growth hormone polyA signal (HGH1) (upper panel) PCR product validation of OGA transgene expression in 2 founder pups (OGA transgenic mice, OGA-TG). The line with the higher OGA expression was chosen for further experiments (lower panel). (D) Western blot for OGA transgene and HA epitope expression in heart [H], gastrocnemius muscle [G], liver [L] and kidney [K] from WT and OGA-TG mice. (E) OGA-TG mice were protected from enhanced AF in T1D. (F) OGA-TG mice had similar blood glucose levels as WT littermates under T1D diabetic and non-diabetic conditions. (G) OGA-TG were protected from enhanced AF in T2D. (H) OGA-TG mice had similar blood glucose levels as WT T2D mice. DON, diazo-5-oxonorleucine; ns, non-significant. Data are represented as mean ± s.e.m, unless otherwise noted. The numerals in the bars represent the sample size in each group. Statistical comparisons were performed using two-tailed Student’s t test or one way AVOVA with Tukey’s multiple comparisons test for continuous variables and Fischer’s exact test for dichotomous variables. (*p <0.05).

We recognized that the results showing AF suppression by DON needed to be interpreted with caution because DON has the potential for off-target effects (76). We thus developed a transgenic mouse model with myocardial overexpression of OGA (OGA-TG mice) (Fig 5C and 5D), as an orthogonal approach to testing for the therapeutic potential of reducing myocardial OGN in diabetic AF. The OGA-TG mice were born in Mendelian ratios, had normal morphology and similar body weight and blood glucose at baseline compared to WT littermates (Supplementary Fig 6A and 6B). These mice had normal echocardiographic features including LV fractional shortening, LV mass and LV dimensions (Supplementary Fig 6C - H) and similar resting heart rates (Supplementary Fig 6I) and heart weight indexed for body weight (Supplementary Fig 6J) compared to WT littermates. The OGA-TG mice were protected from increased AF in T1D compared to WT T1D mice (Fig 5E) and had similar levels of hyperglycemia as WT T1D mice (Fig 5F). Similarly, OGA-TG mice were protected from increased AF in T2D compared to WT T2D mice (Fig 5G) with similar levels of hyperglycemia as the WT T2D mice (Fig 5H). We interpreted these data to show that OGN signaling likely contributes to AF in T1D and T2D.

### Ox-CaMKII and OGN enhance triggered activity and RyR2 Ca^2+^ leak

CaMKII is proarrhythmic, in part, by promoting afterdepolarizations (37, 38, 77, 78). Delayed afterdepolarizations (DADs) are observed as membrane potential fluctuations that occur after action potential repolarization (79). In order to test if DADs were increased in atrial myocytes from T1D mice, we rapidly paced isolated atrial myocytes to simulate the model of AF induction by rapid atrial pacing, using patch clamp in current clamp mode (Fig 6A). Atrial myocytes isolated from WT mice with T1D exhibited increased frequency of DADs and spontaneous action potentials compared to non-diabetic controls (Fig 6B). DON reduced DADs and spontaneous action potentials recorded from atrial myocytes from WT mice with T1D. MMVV mice with T1D were resistant to increased DADs and action potentials, and addition of DON further suppressed DADs and action potentials in these cells (Fig 6B). Taken together, we interpret these data to indicate that ox-CaMKII and OGN augment DADs in T1D, a potential cellular mechanism explaining increased propensity for AF in T1D.

**Figure 6.**
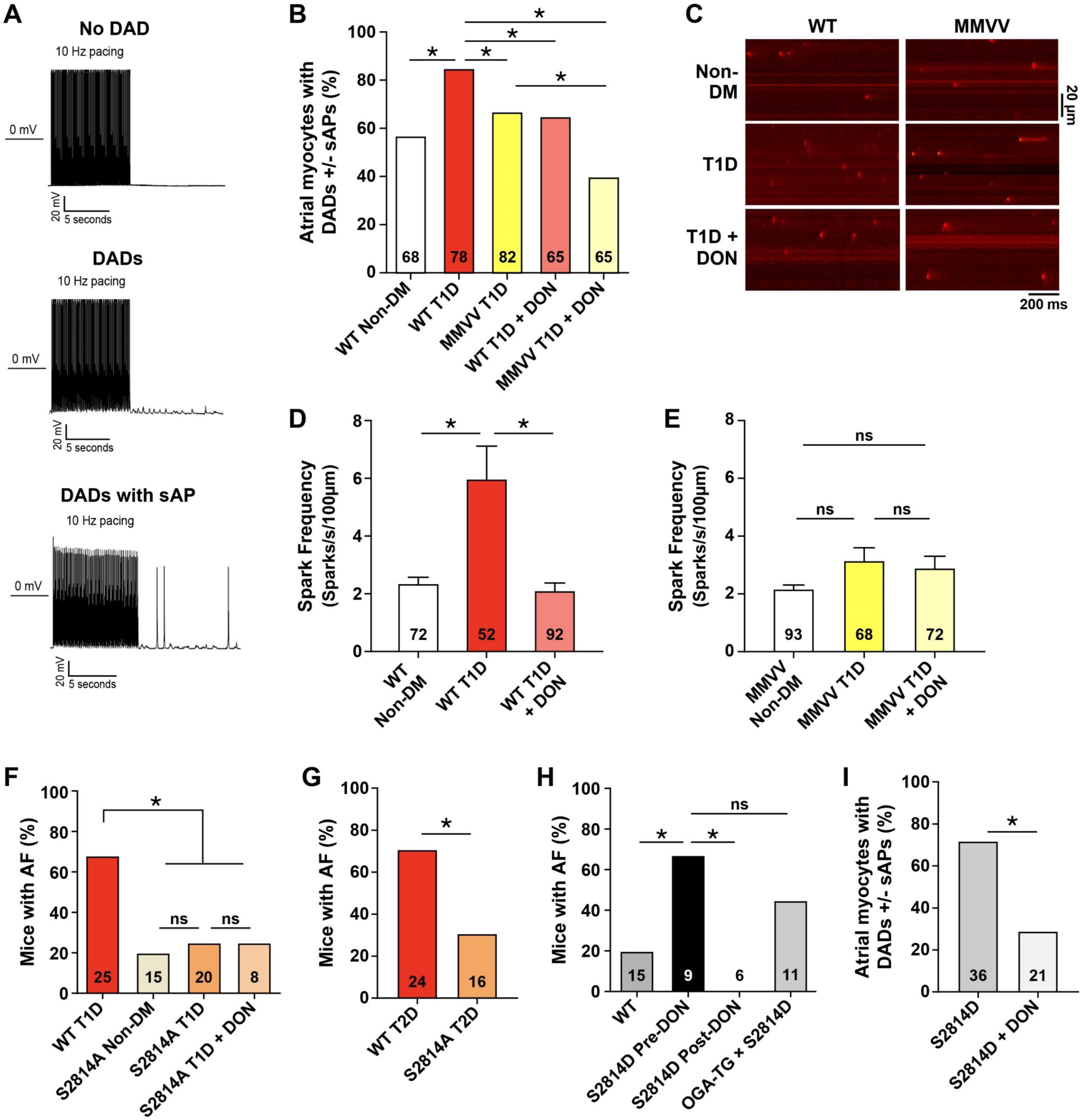
OGA, DON, ox-CaMKII and RyR2 contribute to a proarrhythmic pathway for AF in T1D. (A) Representative images of delayed after depolarizations (DADs) and spontaneous action potentials (sAP) in response to rapid pacing (10Hz) in isolated atrial myocytes. (B) Summary data for combined frequency of DADs and sAPs in response to rapid pacing of isolated atrial myocytes (n = 4 – 7 mice/group). (C) Representative confocal images of Ca^2+^ sparks in atrial myocytes from non-diabetic (non-DM), T1D diabetic and DON pretreated T1D diabetic WT and MMVV mice. Summary data for Ca^2+^ sparks in atrial myocytes from non-DM, T1D and DON pretreated T1D (D) WT and (E) MMVV mice. (n = 52 – 93 cells in each group from 5 – 7 mice/group). S2814A T1D (F) and T2D (G) mice were similarly protected from AF with no further protection in DON pretreated S2814A T1D mice. (H) Summary data of AF incidence in non-diabetic WT, S2814D at baseline (S2814D pre-DON) and S2814D mice treated with DON (S2814D post-DON) after initial AF inducibility at baseline. DON prevented AF in S2814D mice. OGA x S2814D mice were not protected from AF. (I) Summary data for combined frequency of DADs and sAPs in response to rapid pacing of isolated atrial myocytes from S2814D and S2814D mice treated with DON (n = 3 mice/group). Numbers show total number of cells studied per group. Data are means ± s.e.m. Statistical comparisons were performed using two-tailed Student’s t test or one way AVOVA with Tukey’s multiple comparisons test for continuous variables and Fischer’s exact test for dichotomous variables. (*p < 0.05, ns – not significant).

**Figure 7.**
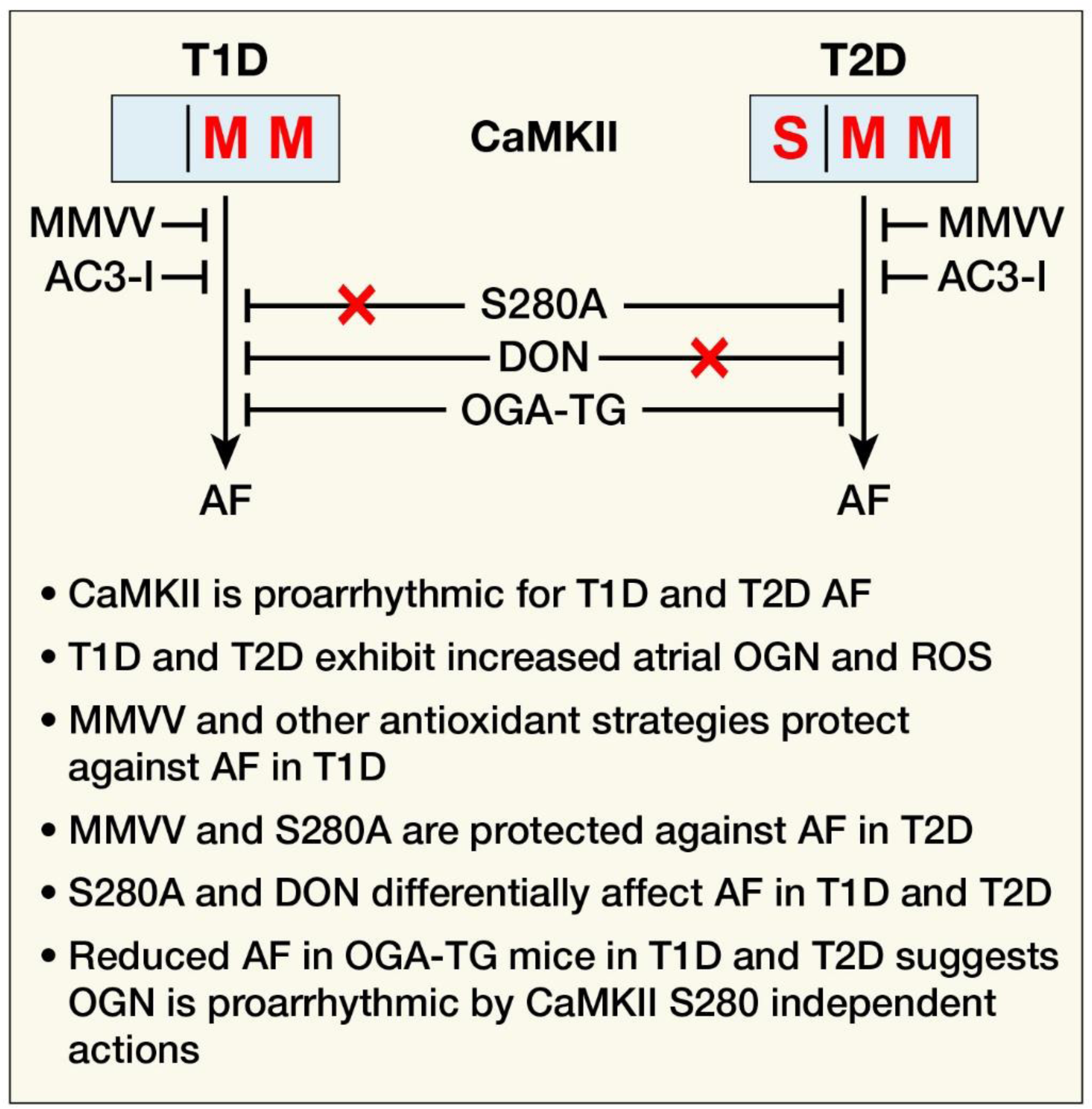
Proposed mechanism for ROS and OGN contributing to AF in diabetes. Signaling through oxidized CaMKII is proarrhythmic in T1D and T2D, while signaling through OGN-CaMKII selectively contributes to proarrhythmia in T2D. DON is preferentially antiarrhythmic in T1D independent of CaMKII. Myocardial OGA overexpression protects from AF in T1D and T2D and OGN is proarrhythmic independent of CaMKII S280. AC3-I mice with genetic myocardial CaMKII inhibition, due to transgenic expression of an inhibitory peptide; DON - Diazo-5- oxonorleucine; MMVV mice are resistant to oxidative CaMKII activation (ox-CaMKII), due to a knock-in replacement of methionines 281/282 with valine in CaMKIIδ; OGA-TG, mice with myocardial OGA overexpression; S280A mice lack an OGN activating CaMKIIδ site, due to a knock-in replacement of serine 280 with alanine in CaMKIIδ.

Excessive RyR2 Ca^2+^ leak has emerged as a core proarrhythmic mechanism that is activated by CaMKII (80). Furthermore, RyR2 is a candidate arrhythmia target downstream to OGN and ox-CaMKII because DON reduced RyR2 Ca^2+^ leak in ventricular cardiomyocytes exposed to hyperglycemia, and suppressed arrhythmias in diabetic rats (31), and MMVV mouse atrial cardiomyocytes were resistant to angiotensin II triggered RyR2 Ca^2+^ leak (32). RyR2 S2814 is a validated CaMKII site (80) and CaMKII catalyzed hyperphosphorylation of S2814 contributes to increased RyR2 Ca^2+^ leak, DADs, and triggered arrhythmias (37, 38, 77, 78). Furthermore, S2814A knock-in mice are resistant to angiotensin II primed AF, a model where ox-CaMKII and RyR2 S2814 are required for proarrhythmia (32). We found increased Ca^2+^ sparks (Fig 6C) in atrial myocytes isolated from WT T1D mice (Fig 6C and 6D), but not from MMVV mice (Fig 6C and 6E). The T1D increased Ca^2+^ spark frequency was suppressed by DON in WT atrial myocytes (Fig 6C and 6D). However, DON did not change the Ca^2+^ spark frequency in MMVV atrial myocytes (Fig 6C and 6E). We found that the S2814A mice were protected from AF in both T1D and T2D (Fig 6F and 6G). We used a knock-in RyR2 S2814 phosphomimetic mutation (S2814D) to test if the antiarrhythmic actions of DON observed in T1D mice required this site (81). We found that AF (Fig 6H) and DADs (Fig 6I) were readily inducible in non-diabetic S2814D mice, and that DON treatment effectively suppressed AF (Fig 6H) and DADs (Fig 6I). In order to directly assess the effect of OGN signaling at the S2814 RYR2 site, without potential off target actions of DON, we interbred the OGA-TG mice with the S2814D mice (OGA-TG x S2814D mice). Surprisingly, the OGA-TG x S2814D mice were susceptible to inducible AF (Fig 6H). We interpreted these data as consistent with the concept that T1D and T2D augment a CaMKII- RyR2 proarrhythmic signaling pathway, but that DON off target actions were antiarrhythmic and/or that the proarrhythmic competence of the CaMKII-RyR2 pathway could be modified by OGN at yet unknown sites. It is possible that the S2814 site undergoes competitive modification by phosphorylation and OGN, and this may explain the lack of AF rescue observed in the OGA- TG x S2814D mice.

Although our data indicates AF is increased in diabetic mice by a pathway that enhances cellular triggering, AF can also be supported by structural tissue level changes, such as cell death and fibrosis, which favor formation of electrical reentrant circuits (77, 82, 83). Furthermore, excessive CaMKII activity can contribute to myocardial death and fibrosis (71, 84, 85). We performed histological analysis of atrial tissue from WT mice with and without T1D (Supplementary Fig 7A and 7B), but did not detect differences in hematoxylin and eosin staining nor fibrosis. We measured cleaved caspase 3, a marker of apoptosis (86), and did not detect differences between WT non-diabetic and WT T1D mice, whereas cleaved caspase 3 was readily detectable in positive control Jurkat cells treated with cytochrome C (Supplementary Figure 7C). We interpret these data to suggest that differences in pathological atrial remodeling were unlikely to contribute to AF in our model, most likely because of the brief, two-week duration of T1D in this model.

## Discussion

### Differential contribution of S280 and M281/282 to AF in T1D and T2D

Our studies provide new information to understand a mechanism for increased AF in DM. We found increased AF in T1D and T2D mouse models, and that this increased susceptibility to AF was absent in mice with myocardial CaMKII inhibition. Thus, CaMKII appears to be a node connecting upstream signals in T1D and T2D with downstream arrhythmia mechanisms. CaMKII is activated by ROS and OGN, and we confirmed previous observations that ROS and OGN are increased in T1D and T2D. However, we found that OGN contributed to AF in T1D and T2D in different ways. Our studies anticipate the need to perform detailed proteomic studies to compare differences in pathways affected by OGN in T1D and T2D. To our knowledge, there are no in vivo data showing CaMKII modification by OGN at S280 or other sites. We were unable to detect this modification on CaMKII from heart lysates isolated from mice with or without T1D and T2D, but did detect phosphorylation at S280 in CaMKIIγ and CaMKIIδ. At this point, it is uncertain if S280 phosphorylation can confer Ca^2+^ and calmodulin-autonomous activity on CaMKII, so future studies will be necessary to test for this possibility. We found that loss of CaMKIIδ S280, the OGN site known to activate CaMKII (31), was insufficient to prevent AF in T1D, whereas loss of this site partially protected against AF susceptibility in T2D.

However, DON, a competitive inhibitor of the hexosamine biosynthetic pathway that reduces availability of UDP-GlcNAc, the rate limiting substrate for OGN, was effective at suppressing AF in T1D but not T2D, suggesting that OGN was proarrhythmic in T1D by targeting sites other than CaMKII, and/or that off target actions of DON were antiarrhythmic. In contrast to our observations in DON treated mice, our finding that OGA TG mice were protected from AF in both T1D and T2D, together with the finding that RyR S2814D knock-in mice were protected from AF by DON but not by interbreeding with the OGA TG mice, suggests that DON is antiarrhythmic, at least in part, by off target actions that are independent of CaMKII, OGN and RyR2 S2814. We interpret the shared resistance to AF by AC3-I, MMVV and OGA-TG mice in T2D and partial protection in S280A mice to suggest that CaMKIIδ and OGN are proarrhythmic in T2D.

In contrast to the different responses of AF in T1D and T2D to manipulation of the OGN pathway, the response to loss of ox-CaMKIIδ was remarkably consistent. The increased susceptibility to AF was similarly suppressed in T1D and T2D in MMVV mice. Upstream of CaMKIIδ, NADPH oxidase and mitochondrial ROS are major sources of intracellular ROS in physiology and disease (87). In this study, we observed protection from AF in T1D in *p47*^-/-^ mice deficient in NADPH oxidase, similar to our previous finding that preventing oxidative activation of CaMKIIδ downstream of NADPH oxidase was antiarrhythmic in an angiotensin II primed model of AF (32). Further, targeted inhibition of mitochondrial ROS with MitoTempo also protected against AF in T1D. Inhibition of mitochondrial ROS by genetic overexpression of the human catalase gene targeted to the mitochondria has been shown to decrease AF in a mouse model (88). The observed rescue with either inhibition of NADPH oxidase or mitochondrial ROS is an interesting finding, and is consistent with the idea of crosstalk between intracellular and mitochondrial ROS production in the feed-forward mechanism known as ROS induced ROS production (89). Although our study is focused on the role of CaMKII, we acknowledge that there are other intracellular targets of oxidation that may play a role in the mechanism of oxidative stress related AF. Indeed, Xie et al. demonstrated that RyR2 oxidation contributes to the pathogenesis of AF (88). We interpret these findings to support an important role for CaMKII as a transducer of pathological ROS in AF in the setting of T1D and T2D. Although we only tested the role of candidate ROS sources, and MsrA in T1D, it seems plausible that future studies will confirm that targeted antioxidant interventions, upstream to CaMKII, have the potential to reduce AF risk in T1D and T2D.

### OGN is a candidate pathway for antiarrhythmic therapy

At present there are no CaMKII-based therapies available for clinical use, but there is evidence that OGN-based therapeutics are advancing to the clinical setting (90–92). The potential that such agents may soon become available is exciting, in part, because they may have broad application, including as antiarrhythmic drugs for AF in patients suffering from DM. If our findings that OGA-TG mice are largely resistant to AF in T1D and T2D have translational implications for humans, then a possible advantage of OGN-based therapies is that they may be successful in T1D and T2D, likely impacting a diverse array of protein targets. While CaMKII and RyR2 appear to be important targets for pathological actions of OGN, OGN targets myriad proteins and pathways. Transcription factors (26), myofilament proteins (30, 93), and mitochondrial metabolism (94), are affected by OGN, and implicated in myocardial disease. OGN is known to participate in physiological processes (26), and is an essential stress response. Mice lacking OGT show augmented pathological responses to cardiac injury (95). Thus, the breadth of OGN substrates, and the requirement for increased OGN to overcome myocardial injury should strike a cautionary note concerning the potential for unintended, off and on target actions with OGN based therapeutics. Availability of drugs, or tool molecules with drug-like properties, will constitute an important step in determining when, how and if OGN therapeutics will be applicable to diabetic patients at risk for AF.

Clinically, the sodium-glucose co-transporter 2 (SGLT2) inhibitors such as empagliflozin and dapagliflozin which were developed primarily as glucose lowering agents in the treatment of T2D, have shown impressive cardiovascular benefits including mortality and heart failure benefits (96, 97). The DAPA-HF clinical trial, showed that these benefits are independent of the presence or absence of diabetes (97). This has led to increased interest in the mechanisms underlying these observed beneficial cardiovascular effects especially as SGLT2 is not expressed in the heart (98, 99). A recent subgroup analysis of the DECLARE-TIMI 58 clinical trial showed that dapagliflozin decreased the incidence of atrial arrhythmias (atrial fibrillation and atrial flutter) in high-risk patients with T2D, independent of HbA1c levels (100). This is in contrast to the other recent class of glucose lowering medications – glucagon-like peptide-1 (GLP-1) agonists, which do not show a benefit in risk reduction of AF despite other cardiovascular benefits (101, 102). In rodent models, it was recently shown that dapagliflozin use in diabetic mice resulted in reduction in myocardial OGN levels and subsequent improvement in ejection fraction in treated compared to untreated mice (103). A recent study showed that empagliflozin attenuated atrial remodeling and improved mitochondrial function in a rat model of T2D (104). Taken together, these findings highlight a potential mechanism and therapeutic target focused on OGN by which SGLT2 inhibitors and especially dapagliflozin can be a tool for the management of AF independent of the presence or absence of diabetes. Exciting early work by Mustroph et al. demonstrates that empagliflozin reduces CaMKII activity and CaMKII-dependent SR Ca^2+^ leak in isolated ventricular myocytes from a mouse heart failure model and human failing ventricular myocytes (99). However the mechanisms by which SGLT2 acts through or on CaMKII are unknown and further work to investigate these mechanisms and their potential antiarrhythmic effects are needed (105).

### Implications for T1D and T2D

T1D represents approximately 10% of all DM (51), and is defined by core features of insulin deficiency and hyperglycemia. In contrast, T2D is the major type of DM, and is mechanistically more complex and heterogenous than T1D. The major risk factor for T2D is obesity, explaining the increased prevalence of T2D worldwide, and the key characteristics of T2D are insulin resistance and hyperglycemia. We chose validated models of T1D (45, 106) and T2D (107, 108) to test the propensity for AF induction in mice. Our findings add new information by which to compare and contrast the consequences of T1D and T2D in heart. While both models relied on an ox-CaMKII pathway for increased AF, and both models showed increased myocardial OGN, T1D and T2D evidenced differential responses to interventions targeting OGN. These differences have the potential to serve as a starting point for future research aiming to interrogate OGN targets, and activity and targeting of OGA and OGT in T1D and T2D.

The shared antiarrhythmic response to loss of CaMKIIδ M281/282, in MMVV knock-in mice, in T1D and T2D coupled with the disparate protection against AF in T2D, but not in T1D by loss of S280, in S280A knock-in mice, has implications for how ROS and OGN are likely to activate CaMKII. Because OGN and ROS are elevated in T1D and T2D, it is possible that CaMKII oxidation and OGN occur together as a hybrid, activating post-translational modification. However, our findings make this seem less probable given the lack of antiarrhythmic response in T1D S280A mice, but shared antiarrhythmic response measured in T1D and T2D MMVV mice. In part because we did not detect OGN at CaMKIIδ S280, we currently lack proteomic information to rule in or rule out the possibility that loss of M281/282 impairs S280 OGN, or, reciprocally, that loss of S280 affects the redox status of M281/282. However, these scenarios would appear more plausible if there were conserved protection or concordant lack of protection against AF in MMVV and S280A mice with T1D or T2D. Although important open questions remain, our data identify CaMKII as a unifying cause of AF in diabetes mellitus. These findings, taken together with emergent evidence that CaMKII contributes to multiple aspects of DM, including gluconeogenesis in liver and insulin signaling in skeletal muscle (109–111), suggest that further study of the role of CaMKII in metabolism and metabolic diseases will be important for improved understanding and treatments.

## Methods

### Mouse models

Experimental studies were performed on male mice with C57BL/6J background. C57BL/6J and mice lacking a functional NADPH oxidase (p47^-/-^) were purchased from The Jackson Laboratory. Our lab previously described the generation of AC3-I (59) and MsrA (71) transgenic mice. We also previously generated MMVV knock-in mice as previously described (46). S2814A and S2814D knock-in mice were generated in the Wehrens laboratory (37, 81). CaMKIIδ-S280A knock-in mice harboring a point mutation in the mouse CaMKIIδ gene to substitute Serine 280 with Alanine (S280A) were generated on a C57BL/6J background using the CRISPR/Cas9 technology. A single-guide RNA (target sequence: 5’-CTGTTGCCTCCATGATGCACAGG-3’) was designed to target CaMKIIδ. Synthetic single-stranded DNA for CRISPR-homology repair was designed to harbor mutations including S280A (TCC --> GCG) and NsiI recognition site (ATGCAT). Genotyping of founder mice and generations of offspring was performed initially by both direct sequencing of PCR amplified fragments and PCR genotyping from tail DNA with the following primers: Forward, 5’-AGGAAATGCTTGCCAAAGTAGTG-3’; Reverse, 5’- CCAGCACATACTGCCCTAGC-3’.

Transgenic mice with myocardial targeted overexpression of OGA (Tg OGA mice) were generated on a C57Bl/6J background. Hemagglutinin (HA)-tagged human Meningioma Expressed Antigen 5 (OGlcNAcase, OGA) cDNA (from the Hart lab) was subcloned into the pBS- αMHC-script-hGH vector between the murine αMHC promoter and a human growth hormone polyadeneylation sequences. Purified pronuclear injections of linearized DNA (digested with NotI) were performed in the Johns Hopkins Transgenic Mouse Core Facility and embryos implanted into pseudo-pregnant females to generate C57Bl6/J F1 mice. The F1 pups were screened for insertion of the transgene into the mouse genome by PCR analysis (Figure 5C), using the forward primer, 5’- TGGTCAGGATCTCTAGATTGGT-3’ and reverse primer, 5’- TCATAAGTTGCTCAGCTTCCTC-3’, producing a product of 850 base pairs. OGA-TG mice were crossed with S2814D mice to generate the OGA-TG x S2814D mice.

### Diabetic mouse models

Type-1 diabetes (T1D) was induced by a single intraperitoneal (i.p.) injection of streptozotocin (STZ) (185mg/kg, Sigma-Aldrich). After a six hour fast, adult (2 – 4 month old) male mice received an i.p. injection of STZ dissolved in a citrate buffer (citric acid and sodium citrate, pH 4.0) or citrate buffer alone for control mice. These mice were maintained on normal chow diet (NCD) (7913 irradiated NIH-31 modified 6% mouse/rat diet – 15 kg, Envigo, Indianapolis, IN) Blood glucose was checked 7 days later with a glucometer (TRUEresult, Nipro Diagnostics) via tail vein and mice with blood glucose levels ≥ 300mg/dl were considered diabetic. STZ-treated mice that had blood glucose levels < 300mg/dl received a second i.p. injection of STZ after a six-hour fast and blood glucose was checked again 7 days after the repeat injection. If blood glucose levels at repeat check was ≥ 300mg/dl, the mice were considered diabetic otherwise they were excluded from the study. Blood glucose, echocardiography and electrophysiology study were done 2 weeks after initiation of STZ injections.

To induce a model of type-2 diabetes (T2D), male mice (C57BL/6J and other mouse genotypes used in this study) aged 5 – 6 weeks were maintained on a high-fat diet (HFD) (Rodent diet with 60 kcal% fat – D12492, Research Diets, New Brunswick, NJ). Five weeks following the initiation of HFD, the mice received daily intraperitoneal (i.p.) injections of low dose streptozocin (STZ) (40 mg/kg/day, Sigma-Aldrich) for three consecutive days after a six hour fast on each day. STZ was dissolved in a citrate buffer (citric acid and sodium citrate, pH 4.0). For control mice, age-matched littermates were maintained on NCD (7913 irradiated NIH-31 modified 6% mouse/rat diet – 15 kg, Envigo, Indianapolis, IN) and received daily i.p. injections of citrate buffer for three consecutive days at similar age as the HFD mice. Blood glucose was checked via tail vein using a glucometer (OneTouch Ultra 2 meter) two weeks after STZ or citrate buffer injection. In addition, body weight, insulin tolerance test, glucose tolerance test and echocardiography were performed two weeks after STZ or citrate buffer injections. Insulin resistance was measured using the Homeostatic Model Assessment of Insulin Resistance (HOMA-IR) calculation based on fasting insulin and glucose levels.

### Blood samples for serum chemistries and renal function

To assess for ketoacidosis and renal function, blood samples for serum chemistries – glucose, blood urea nitrogen, creatinine, and carbon dioxide; were obtained 2 weeks after STZ injection in the T1D mice. Blood sample collection was via facial vein puncture and analysis was performed in the mouse phenotyping core.

### Insulin therapy

Exogenous insulin was delivered subcutaneously via sustained release insulin implants – LinBit (LinShin, Toronto, ON, Canada) (112, 113) to T1D mice with blood glucose levels > 300 mg/dl, one week after STZ injection. Two to 4 implants were implanted subcutaneously according to the manufacturer’s instructions based on the body weight of the mice.

### MitoTEMPO and TPP injections

One week after STZ injection, T1D mice received daily i.p. injections (1mg/kg/day) of MitoTEMPO (2-(2,2,6,6-Tetramethylpiperidin-1-oxyl-4-ylamino)-2-oxoethyl) triphenylphosphonium chloride monohydrate, Enzo Life Sciences, Inc., product number: ALX-430-150-M005) or TPP (Formylmethyl)triphenylphosphonium chloride (2-Oxoethyltriphenylphosphonium chloride) for 7 – 10 days.

### DON injections

T1D mice received DON (diazo-5-oxonorleucine, 5 mg/kg, i.p.) (31) dissolved in phosphate-buffered saline 30 minutes – 1 hour prior to induction of anesthesia for electrophysiology study and rapid atrial burst pacing.

### Mouse electrophysiology and AF induction

In vivo electrophysiology (EP) studies were performed as previously reported with some modifications (32, 37, 57) in mice anesthetized with isoflurane (1.5 – 2% for induction and 1 – 2 % for maintenance of anesthesia; Isotec 100 Series Isoflurane Vaporizer; Harvard Apparatus). Mouse core body temperature (monitored by a rectal probe), heart rate and respiratory rate during the procedure were monitored with a heated surgical monitoring system (MouseMonitorTM S, Indus Instruments, USA) and the body temperature was maintained at 37.0 ± 0.5°C. A Millar 1.1F octapolar EP catheter (EPR-800; Millar/ADInstruments, USA) was introduced via the right jugular vein, as previously described (32, 114), into the right atrium and ventricle for recording intracardiac electrograms. A computer-based data acquisition system (Powerlab 16/30; ADInstruments, USA) simultaneously recorded a 3-lead body surface ECG and up to 4 intracardiac bipolar electrograms (Labchart Pro software, version 7, ADInstruments). Right atrial pacing was achieved by delivering 2-ms current pulses by an external stimulator (STG-3008; Multi Channel Systems). AF inducibility was determined by decremental burst pacing. Burst pacing was started at a cycle length of 40ms, decreasing by 2ms every 2 seconds to a cycle length of 8ms. Burst pacing was performed for a total of five times with an interval of one minute after the end of the previous burst protocol or AF termination. AF was defined as the occurrence of rapid and fragmented atrial electrograms with irregular AV-nodal conduction and ventricular rhythm for at least 1 second. If 1 or more bursts (out of 5) were an AF episode, the mouse was deemed inducible for AF, and non-inducible if there were no AF episodes.

### Transthoracic echocardiography

Transthoracic echocardiography was performed in conscious mice using the Sequoia C256 ultrasound system (Malvern, PA) equipped with a 15-MHz linear transducer as previously described (115, 116). Briefly, M-mode echocardiogram was obtained from parasternal views of the left ventricle at the level of the papillary muscles at sweep speed of 200 mm/sec. Left ventricular dimensions were averaged over 3 to 5 beats at physiological heart rates, and left ventricular mass, fractional shortening and ejection fraction were derived. A blinded operator performed image acquisition and analysis.

### Human samples

De-identified right atrial tissue samples used for CaMKII isoform expression were obtained from patients undergoing cardiac surgery without atrial fibrillation or diabetes.

### Immunoblots

Protein immunodetection by standard western blot techniques was performed. Briefly, mice were sacrificed 2 – 4 weeks after STZ or citrate injection, hearts were excised, flash frozen in liquid nitrogen and stored at -80°C. Hearts were lysed with 1% Triton buffer containing protease inhibitor (P8340, Sigma), phosphatase inhibitor (P0044, Sigma), Thiamet G (SML0244, Sigma), and PUGNAc (A7229, Sigma). Protein fractionation was performed using Nupage gels (Invitrogen) and transferred to PVDF membranes (Bio-Rad). Blots were incubated with primary antibodies – CaMKII delta (1:1000, catalog ab181052, Abcam), *O*-GlcNAc (1:1000, custom antibody – RL2 from the Hart and Zachara lab) (117, 118), MGEA5/OGA (1:2000, catalog ab124807, Abcam), HA epitope tag (1:1000, catalog 600-401-384), cleaved caspase-3 (1:1000, catalog 9662, Cell Signaling Technology), GAPDH (1:10,000, catalog 5174, Cell Signaling); and subsequently with the appropriate HRP-conjugated secondary antibodies – anti-rabbit (1:10,000, catalog 656120, Invitrogen) IgG and anti-mouse (1:10,000, catalog A8786, Sigma-Aldrich) IgM antibodies. Chemiluminescence with ECL reagent (Lumi-Light, Roche) was used for detection. Densitometric analysis was performed using NIH Image J software and band intensity was normalized to the entire Coomassie-stained gel or GAPDH.

### Cellular ROS detection

Hearts were removed immediately after sacrifice, cryopreserved and sectioned at 30-μm thickness. As previously described (71), after washing in phosphate-buffered saline, heart tissue was pre-incubated with DHE for 30 minutes at 37°C and washed. Fluorescence was detected with a laser scanning confocal microscope (Zeiss 510, 40x oil immersion lens) with excitation at 488 nm and detection at 585 nm. The same scanning parameters were applied to all samples and image analysis was done using NIH Image J software.

### Mitochondrial ROS detection

Isolated cardiomyocytes were incubated for 30 minutes with 5μM MitoSOX Red (ThermoFischer Scientific) and 100nM Mitotracker Green (ThermoFischer Scientific). After washing the cells, fluorescence was detected with a laser scanning confocal microscope (Zeiss 510, 40x oil immersion lens). MitoSOX Red (excitation at 510 nm and emission at 580 nm), MitoTracker Green (excitation at 490 nm and emission at 516 nm). Fluorescence was assessed from at least 20 cells in the cell suspension from each mouse. The same scanning parameters were applied to all samples and image analysis was done using NIH Image J software.

### Quantitative PCR for CaMKII isoforms

Total RNA from mouse and human atrial tissue was extracted using Trizol reagent (Invitrogen) according to the manufacturer’s instructions. After first strand cDNA synthesis, messenger RNA expression was analyzed by quantitative real-time polymerase chain reaction on a BioRad CFX machine using pre-validated primers with SsoAdvanced Universal SYBR Green Supermix (Bio-Rad). GAPDH gene expression was used as the reference gene and the specificity of the assay was confirmed by melting curve analysis.

### Neonatal mouse cardiomyocyte isolation, culture and transfection

Cardiomyocytes were isolated from hearts excised from neonatal mice (postnatal day 0 – 3), pooled and dissociated using the MACS Neonatal Heart Dissociation Kit (Catalog #130-098-373), Neonatal Cardiomyocyte Isolation kit, mouse (Catalog #130-100-825) and automated gentleMACS Dissociator (Miltenyl Biotec GmbH), according to the manufacturer’s instructions (119, 120). After isolation, neonatal mouse cardiomyocytes were suspended in Medium-199 (5.5 mM glucose) supplemented with 10% fetal bovine serum, 25 mM HEPES, 50 U/ml penicillin-streptomycin, 2 ug/ml vitamin B12, and 0.1 mM MEM nonessential amino acids. The neonatal cardiomyocytes were plated on 20 µg/ml fibronectin-coated 35 mm glass bottom culture dishes with 20 mm micro-wells (Cellvis, catalog # D35-20-1.5-N) at a seeding density of 1.0 x 10^6^ cells/dish and incubated overnight at 37 °C and 5% CO_2_. The media was changed the next day from to 2% fetal bovine serum media (Medium-199 as described above).

### CaMKII Kinase Translocation Reporter Assay

Construction and validation of the CaMKII kinase activity reporter (CaMKII-KTR) was recently described by our lab in RPE-1 cells and skeletal muscle fibers (73), based on the initial work by and collaboration with Regot et al. (72). CaMKII-KTR was transfected (JetPrime Transfection Reagent, PolyPlus, France) into neonatal mouse cardiomyocytes 24 hours post-isolation. After an 18 hour transfection incubation, cells were imaged in M199 cell culture medium with 2% FBS (Thermo Fisher Scientific, 12340030) supplemented for 18 hours through the experiment with low glucose (5.5 mM), high glucose (33 mM), mannitol (22 mM), NAC (2 μM), or AS105 (1 μM). Caffeine (10 mM) was applied acutely over 20 minutes as a positive control. Fluorescent images were collected using an Olympus IX83 microscope equipped with an ORCA Flash 4.0 sCMOS camera and 20X UPLFLN20X NA0.5 objective lens. Cells were maintained at 37 °C in an OkoLabs stage top incubator. Image analyses were carried out in ImageJ (US National Institutes of Health, Bethesda, MD, United States). The cytosolic to nuclear KTR signal ratios were calculated using the mean intensities measured from the nuclei and cytosolic ring (area immediately perinuclear) of individual cells (72, 73).

### In-Gel Proteolysis for mass spectrometry

Proteins in gel bands, amount 1 μg, were reduced with 1.5 mg/mL dithiothreitol (DTT) in 50 μL of 50 mM of tri-ethyl ammonium bicarbonate (TEAB) buffer at 57 °C for 45min, then alkylated with 10 mg/mL iodoacetomide in 50 μL of 50 mM TEAB buffer in the dark at room temperature for 30 minutes. Buffers were removed by two alternating washes of 50 mM TEAB and acetonitrile re-swelled on ice with 12.5 ng/μL trypsin (Promega, catalog # V5111) in 40 μL of 50 mM TEAB, and proteolyzed at 37 °C overnight, as previously described by Shevchenko et al. (121). Peptides were extracted with 50% acetonitrile/0.1% TFA, dried, resuspended in 0.1% TFA and desalted on Oasis u-HLB plates (Waters, Milford, MA). Peptides were eluted from Oasis plates with 60% acetonitrile/0.1% TFA, dried by vacuum centrifugation, then resuspended in 2% acetonitrile/0.1% formic acid. 30% of samples were used to identify proteins by data-dependent liquid chromatography tandem mass spectrometry (LC-MS/MS).

### LC-MS/MS analysis

Desalted tryptic peptides, were analyzed by nano-liquid chromatography tandem mass spectrometry (nano LC-MS/MS) on an Orbitrap-Lumos mass spectrometer (Thermo Fisher Scientific) interfaced with nano-Acquity LC system (Waters, Milford, MA) using a 2% - 90% acetonitrile/0.1% FA gradient over 70 minutes at 300nl/min on 75 um x 150 mm reverse-phase column (ProntoSIL 120-5-C18 H column 3 µm, 120Å (Bischoff, catalog # F185PS030). Eluting peptides were sprayed into the mass spectrometer through 1 µm emitter tip (New Objective, Woburn, MA) at 2.4 kV. Data dependent analysis was performed by monitoring the top 15 precursor ions in survey scans (350-1800 Da m/z) using a 15s dynamic exclusion. Each prescursor ion was individually isolated with 1.2 Da and fragmented (MS/MS) using HCD activation collision energy 30. Precursor and the fragment ions were analyzed in the orbitrap with resolutions at 200 Da of 120,000 and 30,000, automatic gain control (AGC) targets of 3xe^6^ and 1xe^5^, and maximum injection times (IT) of 60 ms and 200 ms, respectively. Targeted LC-MS/MS analysis was performed under the same chromatography and mass spectrometry conditions. However, instead of monitoring the top 15 precursor ions, a list of m/z values of the STVASMMHR peptide with variable methionine oxidation, serine and threonine O-GlcNAcylation or phosphorylation were monitored, isolated and fragmented.

### Mass spectrometry data analysis

Tandem mass spectra were processed by Proteome Discoverer, v.2.3 (ThermoFisher Scientific) using file re-calibration option and analyzed with Mascot, v.2.6.1 (Matrix Science, London, UK) using Percolator (122) for peptide-spectrum matches (PSM) validation. The Mascot database searches were performed using the RefSeq2017_83_Human with the following criteria: trypsin as enzyme, 2 missed cleavage, 4 ppm precursor mass tolerance, 0.01 Da fragment mass tolerance, and the following variable modifications: oxidation on mehtionine, carbamidomethyl on cysteine, deamidation on asparagine or glutamine, phosphorylation or HexNAcylation on serine or threonine. Tandem mass spectra were also analyzed using PEAKS (v10, Bioinformatics Solutions, Inc.) and Byonic (Protein Metrics) against the same RefSeq2017_83_mus_musculus_database, using the same search criteria as with Mascot, except with mass tolerance of 5 ppm and 0.02 Da on precursor and fragment ions, respectively, in PEAKS and mass tolerances of 5 ppm and 20 ppm on precursor and fragment ions, respectively, in Byonic.

### Calcium imaging in isolated atrial myocytes

Calcium imaging of isolated atrial myocytes was performed as described previously (32, 123).

### Isolated atrial myocyte action potential recording

Voltage clamp studies using the perforated patch configuration on isolated atrial myocytes were performed as described previously to record single cell action potentials (32, 124).

### Fibrosis detection and quantification

Masson’s trichrome staining was used to assess for fibrosis in atrial tissue. Images were digitalized and analyzed using Aperio ImageScope software (Aperio Technologies/Leica Biosystems).

### Statistics

All results are expressed as mean ± s.e.m. for continuous variables or percentages for dichotomous variables. For continuous variables, statistical analyses was performed using an unpaired Student’s t-test (2-tailed – for 2 groups) or one-way ANOVA followed by Tukey’s or Dunnetts post-hoc multiple comparisons (for ≥ 3 groups). For dichotomous variables, statistical analyses were performed using Fischer’s exact test (2-tailed for 2 groups) or a Chi-square test with appropriate correction for post-hoc multiple comparisons (for ≥ 3 groups). A p-value of < 0.05 was considered statistically significant.

### Study approval

The Johns Hopkins University animal care and use committee approved all animal experimental protocols. For human samples, the local institutional review board of the Goettingen University approved all human procedures and each patient gave written informed consent.

## Abbreviations

AF: Atrial fibrillation
CaMKII: Calcium and calmodulin-dependent protein kinase II
DM: Diabetes mellitus
DON: Diazo-5-oxonorleucine
KTR: Kinase translocation reporter
MsrA: Methionine sulfoxide reductase A
OGA: O-GlcNAcase
OGN: O-GlcNAcylation
(OGT): O-GlcNAc transferase
OGN-CaMKII: O-GlcNAcylated CaMKII
ox-CaMKII: Oxidized CaMKII
STZ: Streptozocin
T1D: Type 1 diabetes mellitus
T2D: Type 2 diabetes mellitus
RyR2: Type 2 ryanodine receptors

## Author Contributions

MEA, OOM and AGR conceived the project, GWH, PSB, RSA, NEZ, PU and EDL contributed important input for planning the studies, and MEA and OOM wrote the manuscript. NA and JLP developed the S280A mouse. OOM and AT developed the OGA-TG mouse. XHW provided the S2814A and S2814D mice. BC, L-SS, and YW performed cellular electrophysiological and Ca^2+^ measurement studies. LSM and AGR provided human specimens. OOM, JMG, QW, KRM, EDL, NEZ and PSB performed CaMKII assays and OGN studies. OOM, EDL, TNB and RNC performed mass spectrometry studies. OOM and QW performed quantitative PCR studies. All authors edited and approved the final manuscript.

## Acknowledgements

We are grateful to Jinying Yang for her assistance in maintaining mouse colonies and Gianna Bortoli for her assistance with neonatal cell isolation. We are also grateful to Djahida Bedja for performing the mouse echocardiograms. We thank Chip Hawkins and the Johns Hopkins University School of Medicine Transgenic Core Laboratory for their technical expertise in generating transgenic mice. We thank the Johns Hopkins University of School of Medicine Mass Spectrometry and Proteomics Facility for their technical assistance in the mass spectrometry studies. We also thank Teresa Ruggle who produced the artwork. We are grateful to Howard Schulman for the gift of the CaMKII inhibitor - AS105.

---

**Figure S1.**
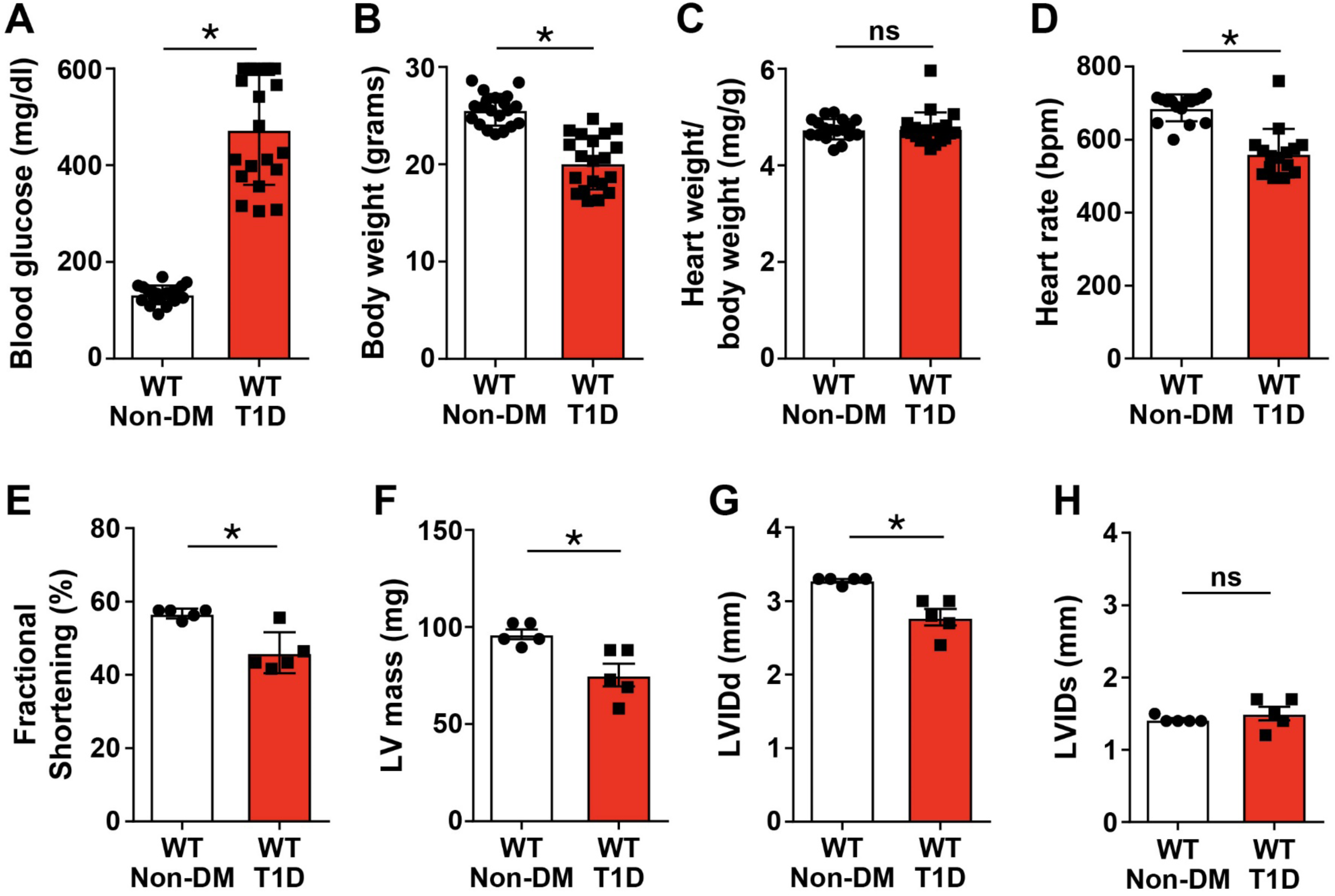
T1D mouse model. Summary data for (A) blood glucose and (B) body weight one week after STZ injection (n = 20 WT non-DM, n = 20 WT T1D); Summary data for (C) heart weight indexed for body weight and (D) heart rate two weeks after STZ injection (n = 18 WT non-DM, n = 20 WT T1D); Summary data of echocardiographic parameters two weeks after STZ injection – fractional shortening (E), LV mass (F), LVIDd (G) and LVIDs (H) in WT non-DM (n = 5) and WT T1D (n = 5) CTRL, citrate buffer; EP, electrophysiology study; LV, left ventricular; LVIDd, LV internal diameter end diastole; LVIDs, LV internal diameter end systole; STZ, streptozocin. Data are represented as mean ± s.e.m., significance was determined using two-tailed Student’s t test. (*p < 0.05, ns – not significant).

**Figure S2.**
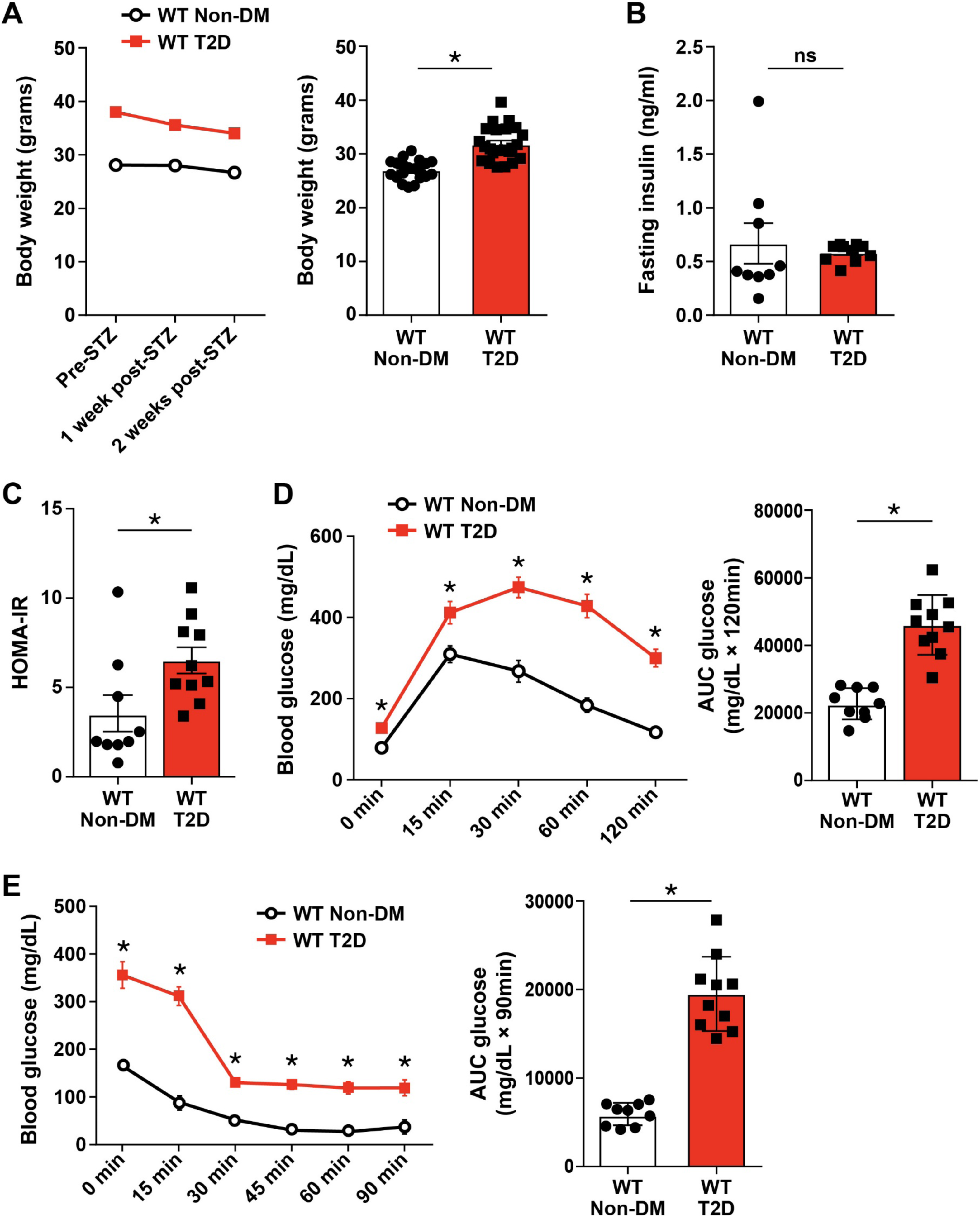
T2D mouse model. Summary data for (A) body weight trend (left panel) after five weeks of high-fat diet (HFD) and subsequent daily doses of low dose STZ for three consecutive days (n = 7 WT non-DM, n = 10 WT T2D), and body weight (right panel) two weeks of after STZ injection at time of EP study (n = 21 WT non-DM, n = 24 WT T2D) (B) Summary data for fasting insulin level in WT non-DM and T2D mice two weeks after STZ injection; T2D mice had features consistent with T2D as determined by homeostatic model assessment of insulin resistance, HOMA-IR (C), glucose tolerance test (D) and insulin tolerance test (E) (n = 9 WT non-DM, n = 10 WT T2D). AUC, area under the curve; GTT, glucose tolerance test; ITT, insulin tolerance test. Data are represented as mean ± s.e.m.; significance was determined using two-tailed Student’s t test. (*p < 0.05, ns – not significant).

**Figure S3.**
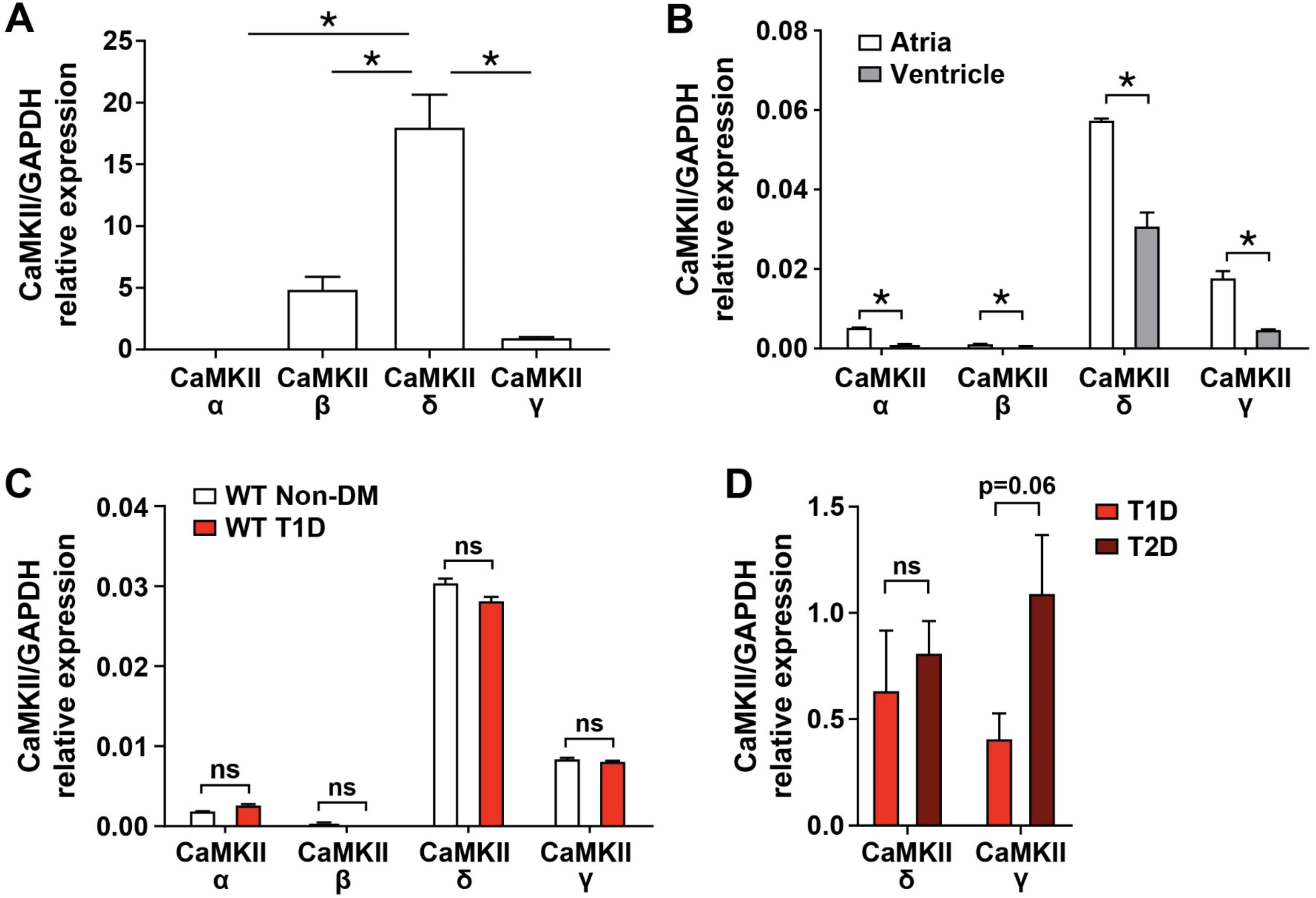
CaMKIIδ is the predominant isoform in mouse and human atria. (A) CaMKII isoforms (α, β, γ, δ) mRNA expression in human atria from non-diabetic patients with no history of AF measured by qRT-PCR using total RNA (n = 5). See Supplemental Table 2 for patient characteristics. (B) CaMKII isoforms (α, β, γ, δ) mRNA expression in atria and ventricles from WT non-diabetic mice measured by qRT-PCR using total RNA (n = 5). (C) CaMKII mRNA expression in atria from non-diabetic and WT T1D mice (n = 5 WT non-DM, n = 5 WT T1D). (D) CaMKIIγ and CaMKIIδ mRNA expression in atria from WT T1D compared with WT T2D mice (n = 4 WT T1D, n = 4 WT T2D). Data are represented as mean ± s.e.m. Statistical comparisons were performed using two-tailed Student’s t test or one way AVOVA with Tukey’s multiple comparison’s post-hoc test. (*p < 0.05, ns – not significant).

**Figure S4.**
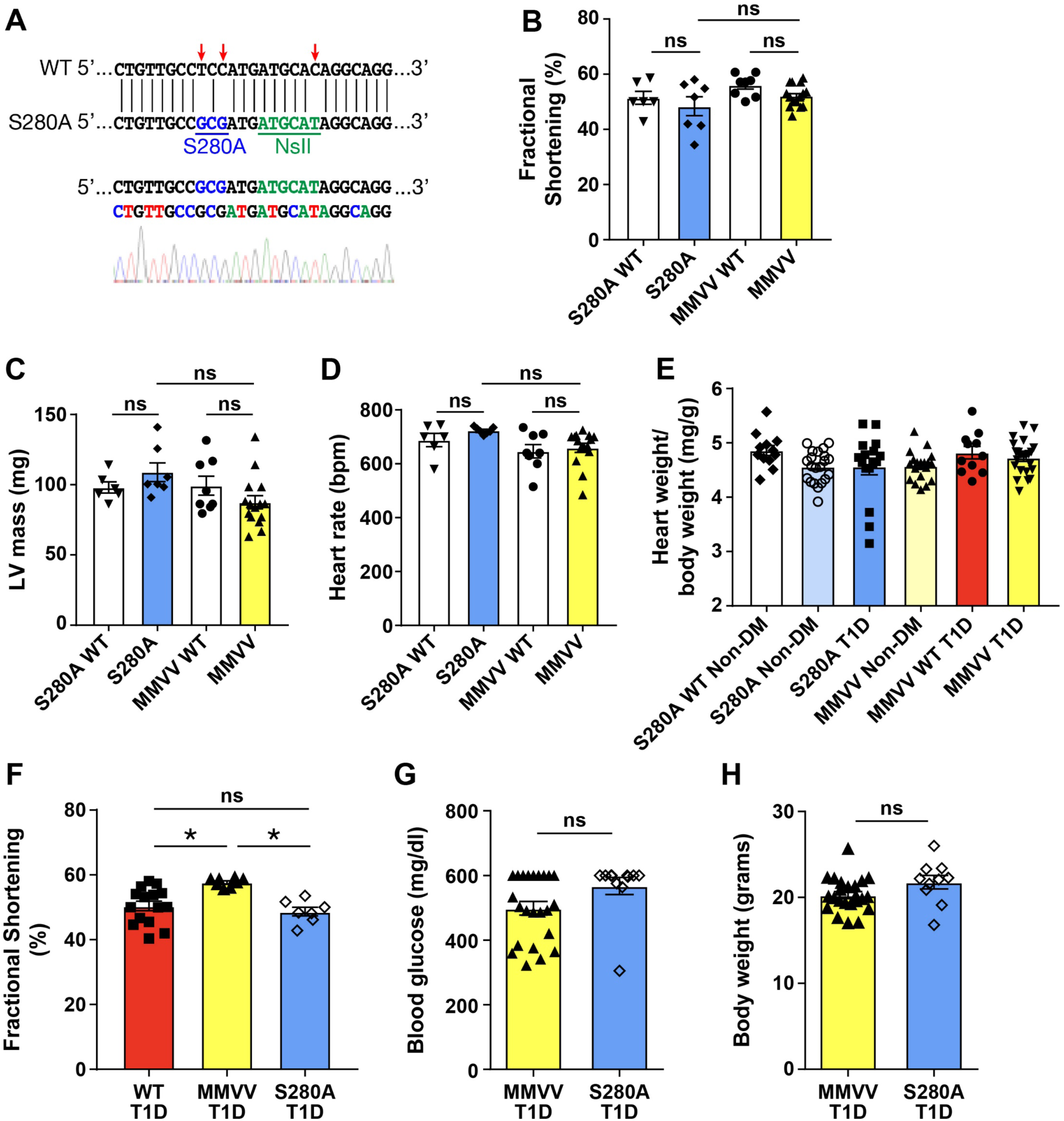
Characteristics of S280A and MMVV mice. (A) Design of knock-in sequence and Nsil genotyping site for S280A mice (upper panel). F2 sequence showing homozygous S280A (lower panel) (B) Summary data for echocardiographic analysis of non-diabetic S280A and MMVV mice revealed no difference in fractional shortening (B), LV mass (C) or heart rate (D) compared to WT littermates (n = 6 S280A WT, n = 7 S280A, n = 8 MMVV WT, n = 15 MMVV). (E) Summary data for heart weight indexed for body weight in non-diabetic and T1D S280A and MMVV mice and WT littermate controls (n = 12 S280A WT non-DM, n = 21 S280A non-DM, n = 16 S280A T1D, n = 19 MMVV non-DM, n = 11 MMVV WT T1D, n = 23 MMVV T1D). (F) Summary data for left ventricular fractional shortening in T1D WT, MMVV and S280A mice. MMVV mice were protected from diabetic cardiomyopathy compared to T1D WT and MMVV mice (n = 23 MMVV T1D, n = 11 S280A T1D). Summary data for blood glucose (G) and body weight (H) in T1D MMVV and S280A mice, two weeks after STZ injection (n = 23 MMVV T1D, n = 10 S280A T1D). Data are represented as mean ± s.e.m., statistical comparisons were performed using two-tailed Student’s t test or one way AVOVA with Tukey’s multiple comparison’s test. (*p <0.05 ns – not significant).

**Figure S5.**
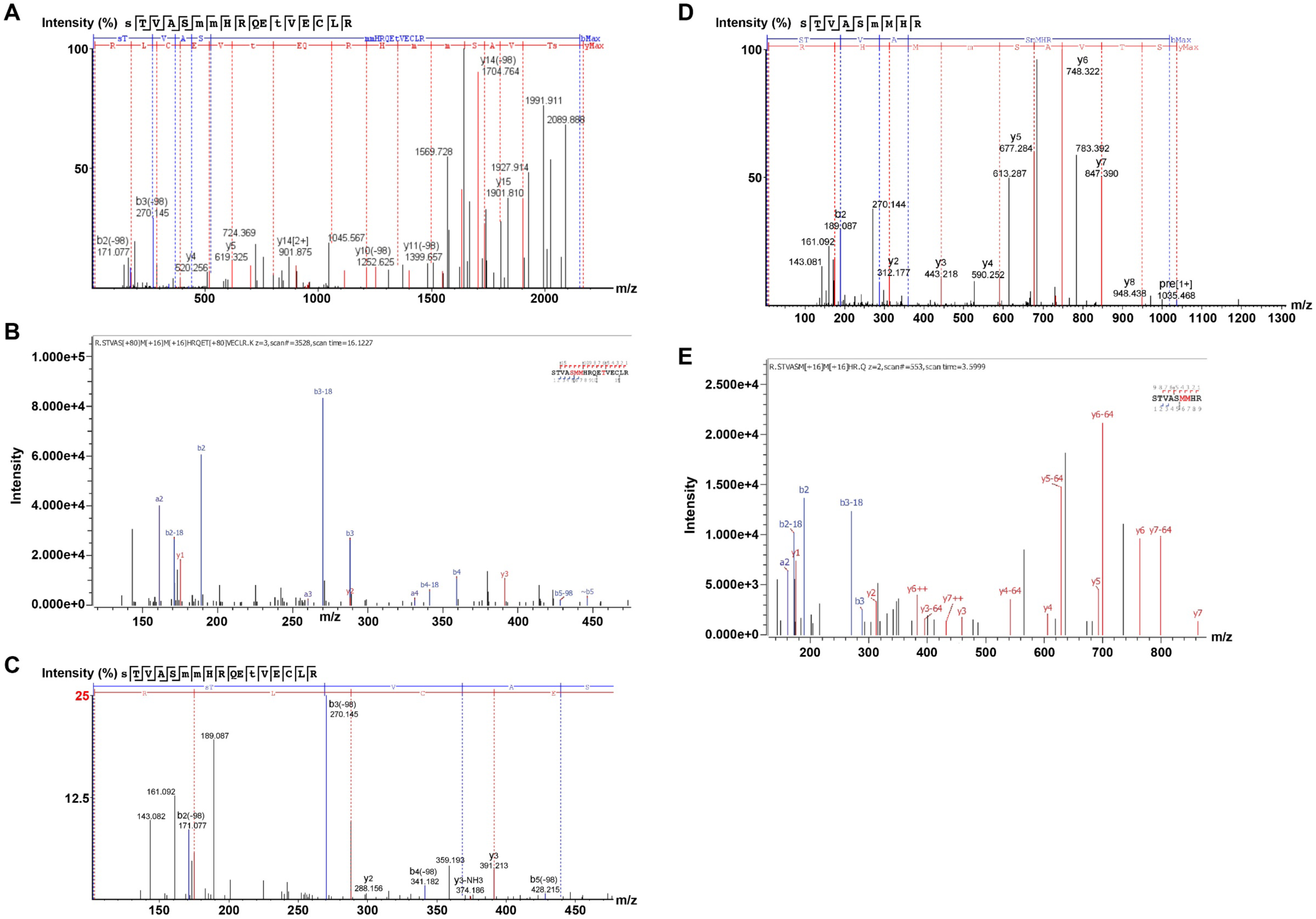
Tandem MS analysis showed phosphorylation but no OGN at S280, and confirmed oxidation at MM281/282 and phosphorylation at T287 in diabetic hearts. Heart lysates from non-diabetic and diabetic hearts were subjected to in-gel proteolysis followed by LC-MS/MS analysis with a focus on the peptide – STVASMMHR. The observed fragment ions are indicated in the sequence for each spectra. Small case letters and red letters indicate fragment ions that contain the mass shift indicative of the respective modifications (A) Fragmentation of di-phosphorylated and di-oxidized STVASMMHRQETVECLR shows phosphorylation at S276 and T287, and oxidization at M281 and M282. A neutral loss of -98 Da (H3PO4) represents loss of the phosphate group plus water from Ser or Thr and the loss of -64 Da represents the loss of the oxidized sulfate group from methionine. (B) Expanded m/z range from spectra in A showing b3 ion which maps phosphorylation to S280 and (C) a 270.145 m/z ion that is either b3-18 (loss of H2O) if phosphate is on S280 or b3-98 (loss of H3PO4) if phosphate is at S276 or T277, and is consistent with a chimera spectrum containing both di-phosphorylated forms of this peptide. Fragmentation spectra show STVASMMHR is oxidized at M281 (D) and di-oxidized at M281 and M282 (E).

**Figure S6.**
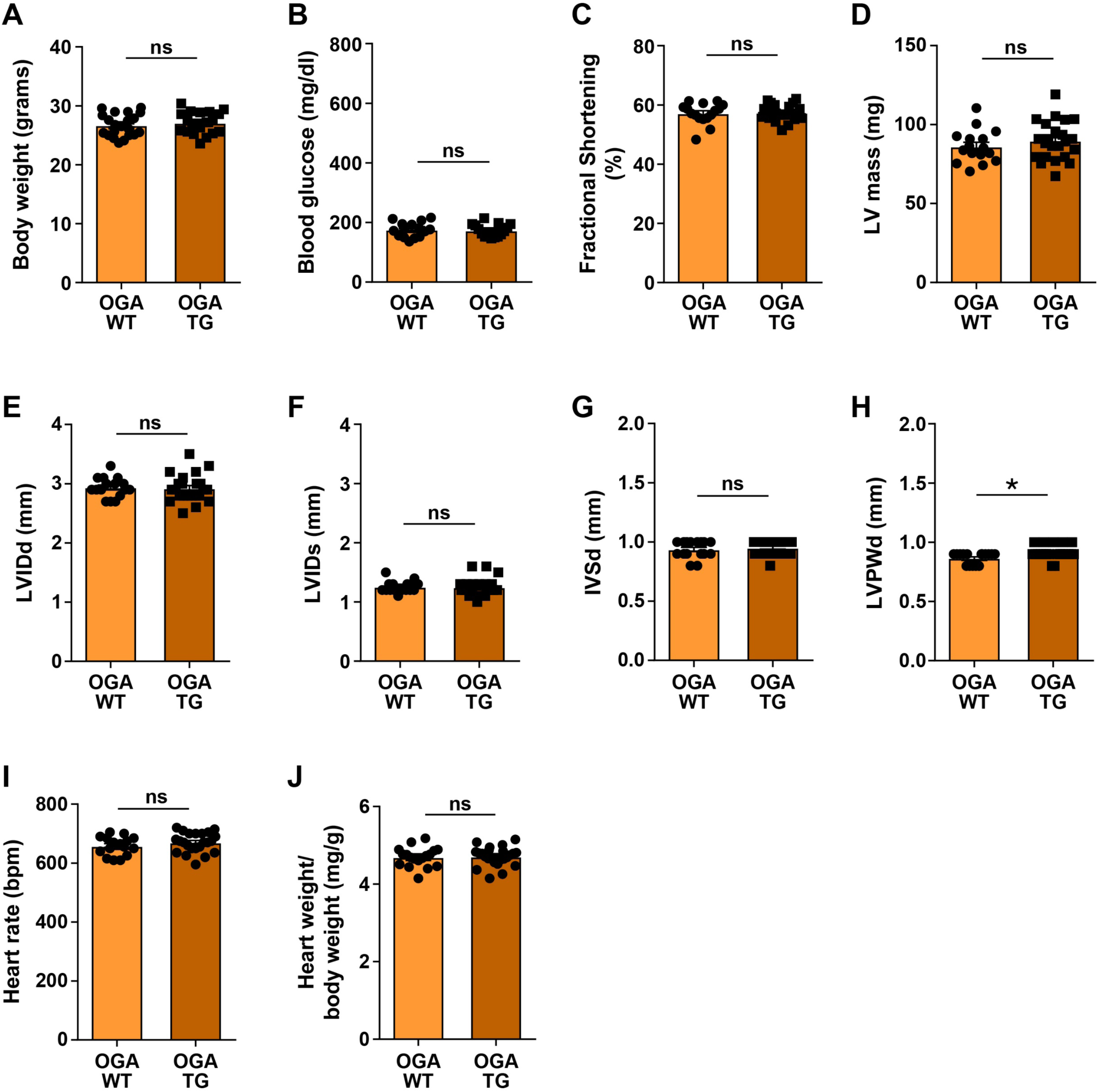
Cardiac phenotyping of OGA-TG mice. (A) Summary data for body weight in WT (n = 21) and OGA-TG (n = 19) mice. (B) Summary data for baseline blood glucose in non-diabetic WT (n = 15) and OGA-TG (n = 15) mice. Summary data of echocardiographic parameters in non-diabetic WT (n = 16) and OGA-TG (n = 23) mice – LV fractional shortening (C), LV mass (D), LVIDd (E), LVIDs (F), IVSd (G), LVPWd (H) and heart rate (I). There were no differences in any of these parameters. (J) Summary data for heart weight indexed for body weight in WT (n = 18) and OGA-TG (n = 24). IVSd, interventricular septal end diastolic dimension; LVPWd, left ventricular posterior wall end diastolic dimension. Data are represented as mean ± s.e.m., statistical comparisons were performed using two-tailed Student’s t test. (*p <0.05, ns – not significant).

**Figure S7.**
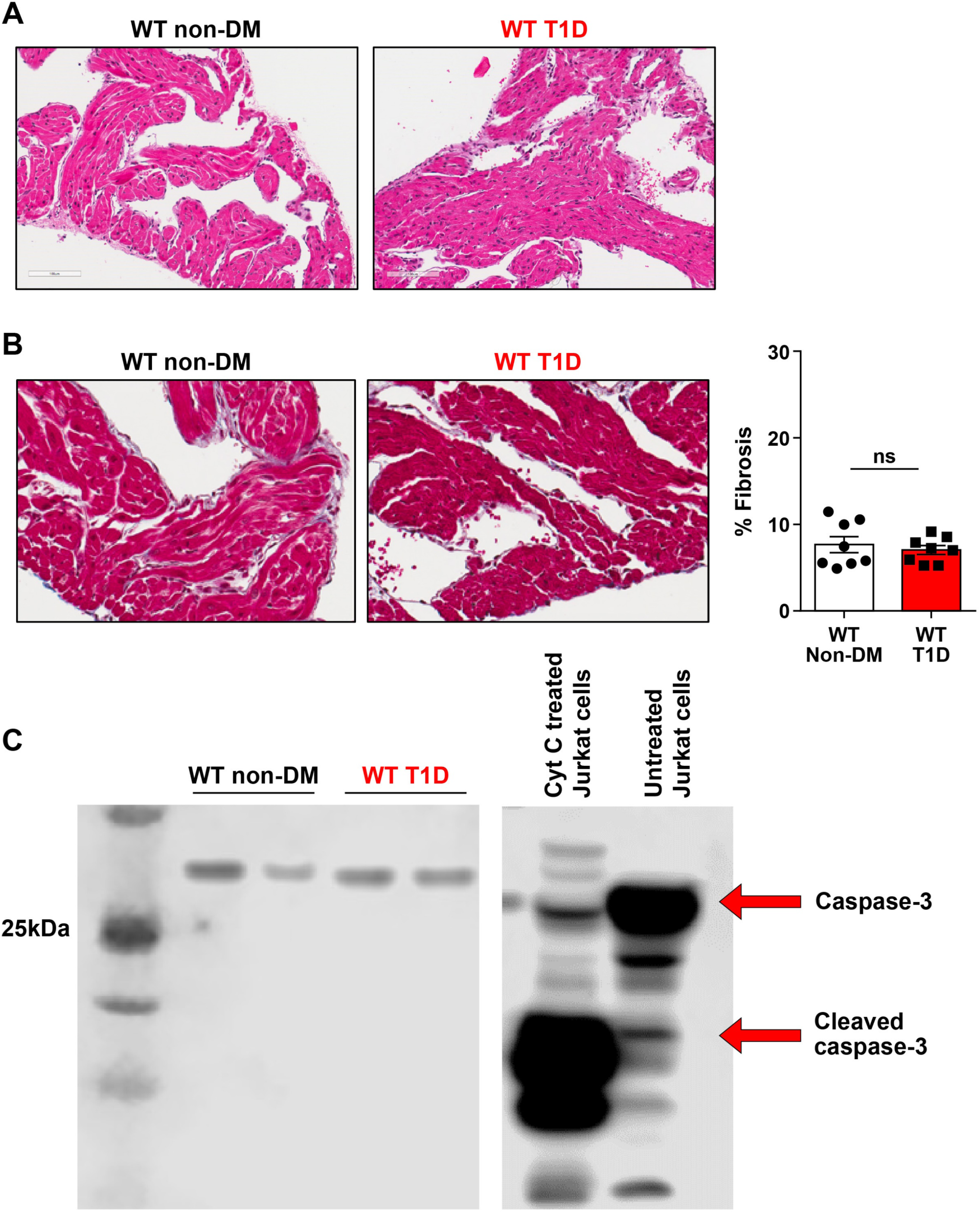
Absence of increased atrial fibrosis or apoptosis in T1D mice. (A) Representative images of hematoxylin and eosin staining of atrial tissue from WT non-DM and WT T1D mice (B) Representative images of Masson’s trichrome staining of atrial tissue from WT non-DM and WT T1D mice. There was no increased fibrosis in the T1D mice compared to non-DM mice. (C) Immunoblots for caspase-3 and cleaved caspase-3 in atrial tissue from WT non-DM and WT T1D mice showed no evidence of increased apoptosis in the T1D mice. Cytochrome C treated and untreated Jurkat cells as positive and negative controls respectively for cleaved caspase-3. Cyt C, cytochrome C. Data are represented as mean ± s.e.m., statistical comparisons were performed using two-tailed Student’s t test. (ns – not significant).

**Supplementary Table 1.** Cardiac morphometric measures and serum chemistry of WT Ctrl and T1D mice at electrophysiology study.

|  | **WT Ctrl** | **WT T1D** |
| --- | --- | --- |
| **Heart weight/tibial length (mg/mm)** | 6.85 ± 0.18, n=6 | 4.93 ± 0.15, n=9* |
| **Atrial weight/tibial length (mg/mm)** | 0.53 ± 0.03, n=6 | 0.38 ± 0.02, n=9* |
| **Ventricular weight/tibial length (mg/mm)** | 6.18 ± 0.20, n=6 | 4.43 ± 0.15, n=9* |
| **CO<sub>2</sub>** | 15.5 ± 1.1 | 16.8 ± 0.7 |
| **BUN (mg/dl)** | 24.6 ± 0.7 | 24.2 ± 1.7 |
| **Cr (mg/dl)** | 0.4 ± 0 | 0.4 ± 0.03 |
| **Serum blood glucose (mg/dl)** | 183.4 ± 13.4 | 697.4 ± 48.42* |

Heart weight normalized to tibial length, atrial weight normalized to tibial length, and ventricular weight normalized to tibial length (\*p < 0.05); n = 6 – 9/group. Serum pCO<sub>2</sub>, BUN, creatinine and blood glucose. (\*p < 0.05, n = 4 – 5/group). Data are means ± s.e.m

**Supplementary Table 2.** Patient characteristics for atrial samples.

| **Patient Characteristics** | **Sinus rhythm and no DM (n = 5)** |
| --- | --- |
| Age (year) | 68 ± 4 |
| Male:Female | 2:3 |
| Ejection Fraction (%) | 57 ± 9 |
| Drug Treatment |  |
| B-blocker, n (%) | 4/5 (80) |
| ACEI/ARB, n (%) | 5/5 (100) |
| Calcium channel blocker, n (%) | 0/5 (0) |
| Statin, n (%) | 5/5 (100) |

ACEI, Angiotensin converting enzyme inhibitor; ARB – Angiotensin receptor blocker.

**Supplementary Table 3.** Coverage of CaMKII isoforms in mass spectrometric analysis of heart lysate samples

| Assession Number | Protein | Percent Coverage | Unique Peptides | O-GlcNAc Sites Identified | Phosphorylation Sites Identified | Engine |
| --- | --- | --- | --- | --- | --- | --- |
| XP_017174906 | CaMKII delta isoform X8 (X7, X12, others) | 57.7 | 4 (64) | no | S276, T277, T287 | Byonic, PEAKS |
| XP_017174906 | CaMKII delta isoform X8 (X7, X12, others) | 57.7 | 4 (64) | no | S276, T277<br>S280, T287 | PD-Mascot-Percolator |
| XP_006518551.1 | CaMKII gamma isoform X1 (X7) | 53.4 | 4 | no | S280, T287 | Byonic (Pep 3e-8) |
| XP_006518557.1 | CaMKII gamma isoform X7 (X6) | 46.2 | 4 (35) | no | S280, T287 | PD-Mascot-Percolator, Byonic (PEP 1.4e-19) |
| XP_006540759.1 | PREDICTED: nucleobindin-1 isoform X1 | 32% | 13 | S38/T42/T44 | none | PD-Mascot-Percolator; Byonic |

**Supplementary Table 4.** Calculated b and y fragment ions of STVASMMHRQETVECLR with phosphorylation on S280 and T287 and oxidation on M281 and M282. Red m/z values are present in Supplementary Figures 5A and 5B within <0.01 m/z.

| # | a calc. | b calc. | b-18 calc. | b++ calc. | Seq | y calc. | y++ calc. | # |
| --- | --- | --- | --- | --- | --- | --- | --- | --- |
| 1 | 60.0444 | 88.0393 | 70.0287 | 44.5233 | S |  |  | 17 |
| 2 | 161.0921 | 189.087 | 171.0764 | 95.0471 | T | 2082.8208 | 1041.914 | 16 |
| 3 | 260.1605 | 288.1554 | 270.1448 | 144.5813 | V | 1981.7731 | 991.3902 | 15 |
| 4 | 331.1976 | 359.1925 | 341.1819 | 180.0999 | A | 1882.7047 | 941.856 | 14 |
| 5 | 498.196 | 526.1909 | 508.1803 | 263.5991 | S(+79.97) | 1811.6676 | 906.3374 | 13 |
| 6 | 645.2314 | 673.2263 | 655.2157 | 337.1168 | M(+ 15.99) | 1644.6692 | 822.8382 | 12 |
| 7 | 792.2668 | 820.2617 | 802.2511 | 410.6345 | M(+ 15.99) | 1497.6338 | 749.3205 | 11 |
| 8 | 929.3257 | 957.3206 | 939.31 | 479.1639 | H | 1350.5984 | 675.8028 | 10 |
| 9 | 1085.427 | 1113.422 | 1095.411 | 557.2145 | R | 1213.5395 | 607.2734 | 9 |
| 10 | 1213.485 | 1241.48 | 1223.47 | 621.2438 | Q | 1057.4384 | 529.2228 | 8 |
| 11 | 1342.528 | 1370.523 | 1352.512 | 685.7651 | E | 929.3798 | 465.1935 | 7 |
| 12 | 1523.542 | 1551.537 | 1533.526 | 776.2721 | T(+ 79.97) | 800.3372 | 400.6722 | 6 |
| 13 | 1622.61 | 1650.605 | 1632.595 | 825.8063 | V | 619.3232 | 310.1652 | 5 |
| 14 | 1751.653 | 1779.648 | 1761.637 | 890.3276 | E | 520.2548 | 260.631 | 4 |
| 15 | 1854.662 | 1882.657 | 1864.647 | 941.8322 | C | 391.2122 | 196.1097 | 3 |
| 16 | 1967.746 | 1995.741 | 1977.731 | 998.3742 | L | 288.203 | 144.6051 | 2 |
| 17 |  |  |  |  | R | 175.119 | 88.0631 | 1 |

**Supplementary Table 5.** Calculated b and y fragment ions of STVASMMHRQETVECLR with phosphorylation on S276 and T287 and oxidation on M281 and M282. Red m/z values are present in Supplementary Figures 5A and 5C within <0.01 m/z.

| # | Immonium | b | b-NH3 | b (2+) | Seq | y | y-NH3 | Y (2+) | # |
| --- | --- | --- | --- | --- | --- | --- | --- | --- | --- |
| 1 | 140.011 | 168.006 | 150.979 | 84.503 | S(+ 79.97) |  |  |  | 17 |
| 2 | 74.061 | 269.054 | 252.027 | 135.027 | T | 2002.854 | 1985.828 | 1001.927 | 16 |
| 3 | 72.081 | 368.122 | 351.095 | 184.561 | V | 1901.810 | 1884.780 | 951.403 | 15 |
| 4 | 44.050 | 439.159 | 422.132 | 220.080 | A | 1802.738 | 1785.711 | 901.875 | 14 |
| 5 | 60.045 | 526.191 | 509.164 | 263.596 | S | 1731.704 | 1714.674 | 866.351 | 13 |
| 6 | 120.048 | 673.227 | 656.200 | 337.113 | M(+ 15.99) | 1644.669 | 1627.642 | 822.835 | 12 |
| 7 | 120.048 | 820.262 | 803.235 | 410.631 | M(+ 15.99) | 1497.634 | 1480.6071 | 749.317 | 11 |
| 8 | 110.072 | 957.321 | 940.294 | 479.161 | H | 1350.598 | 1333.571 | 675.799 | 10 |
| 9 | 129.114 | 1113.422 | 1096.395 | 557.211 | R | 1213.539 | 1196.512 | 607.270 | 9 |
| 10 | 101.071 | 1241.481 | 1224.454 | 621.240 | Q | 1057.438 | 1040.411 | 529.219 | 8 |
| 11 | 102.055 | 1370.523 | 1353.496 | 685.762 | E | 929.380 | 912.353 | 465.190 | 7 |
| 12 | 154.027 | 1551.537 | 1534.510 | 776.269 | T(+ 79.97) | 800.337 | 783.310 | 400.669 | 6 |
| 13 | 72.081 | 1650.606 | 1633.579 | 825.803 | V | 619.325 | 602.296 | 310.162 | 5 |
| 14 | 102.055 | 1779.648 | 1762.621 | 890.324 | E | 520.256 | 503.228 | 260.627 | 4 |
| 15 | 76.022 | 1882.658 | 1865.631 | 941.829 | C | 391.213 | 374.186 | 196.106 | 3 |
| 16 | 86.097 | 1995.742 | 1978.715 | 998.371 | L | 288.203 | 271.176 | 144.602 | 2 |
| 17 | 129.114 |  |  |  | R | 175.119 | 158.092 | 88.059 | 1 |

